# MetaboRamics: Highly multiplexed metabolic imaging by stimulated Raman for spatial metabolomics in live cells

**DOI:** 10.64898/2026.05.21.727012

**Authors:** Rahuljeet S. Chadha, Adrian Colazo, Joseph A. Ambarian, Ziguang Yang, Philip A. Kocheril, Benjamin Yang, Lu Wei

## Abstract

Cellular metabolism is highly dynamic, intertwined and spatially heterogenous, yet methods that can simultaneously visualize multiple metabolic pathways in living systems remain largely limited. Here, we present MetaboRamics: highly multiplexed **metabo**lic imaging by stimulated **Ram**an for spatial metabolom**ics** in live cells. Through rational probe selection and careful optimization, isotope editing, and robust spectral unmixing, we establish a metabolic palette of 16-colors spanning glucose uptake and utilization, lipid uptake and synthesis, choline metabolism, DNA synthesis, and amino acid incorporation in addition to endogenous proteins, lipids and redox signals. Incorporation of organelle-targeted Raman probes further enables spatial interactomics and assessment of organelle activities. Applying MetaboRamics to epithelial-mesenchymal transition (EMT), we observe metabolic rewiring in mesenchymal cells, reflected by reduced glucose-derived biomass, lipid turnover, protein synthesis, and altered redox balance. Finally, we perform optical phenotyping of cellular states under metabolic stress of serum deprivation, nutrient overload (fructose, and saturated fatty acid), inflammation, and pharmacological perturbations to reveal subcellular metabolic changes. This work fully realizes the potential of stimulated Raman scattering (SRS) microscopy for super-multiplexed metabolic imaging by establishing, for the first time, a 16-plex platform for live-cell spatial metabolomics.

## Introduction

Metabolism is an inherently dynamic, spatially organized network of chemical reactions that enables essential cellular functions. Changes in metabolic flux often initiate or accompany disease progression, making metabolism a critical lens for investigating disease. Recent advances in high-throughput genomics,^1,2^ transcriptomics^3,4^ including spatial transcriptomics,^5–7^ and proteomics^8–10^ have transformed our understanding of cellular states by allowing comprehensive, scalable, and increasingly spatially resolved measurements. These advances are largely driven by sequencing-based technologies that leverage molecular amplification and the stability of linear biopolymers such as DNA and RNA. In contrast, metabolites cannot be amplified, are difficult to chemically fix, and exhibit extensive chemical diversity, lability, low abundance, and broad dynamic range.^11^ Consequently, spatial metabolomics remains a grand challenge, and approaches that can directly visualize multiple metabolic activities in living cells are still highly limited, despite the central role of metabolism in health and disease.

Recently, mass spectrometry imaging (MSI), leveraging matrix-assisted laser desorption/ionization (MALDI),^12,13^ secondary ion mass spectrometry (SIMS),^14^ and desorption electrospray ionization (DESI),^15^ have significantly expanded the capability for *in situ* spatial metabolomics. Imaging mass cytometry has also been demonstrated to perform untargeted spatial metabolomics and targeted multiplexed protein imaging using a single workflow.^16^ However, despite achieving excellent molecular sensitivity and specificity, these techniques are inherently destructive and thus incompatible with live-cell subcellular analysis. Moreover, additional sample processing requirements and variable ionization efficiencies pose additional challenges for throughput and quantitative accuracy. Alternatively, fluorescence microscopy enables ultrasensitive live-cell analysis, but relies on bulky fluorophores or genetically encoded reporters that can perturb native biology, are substantially larger than the metabolites of interest, and possess inherently broad spectra, thereby limiting multiplexing for metabolomic applications.^17^

To address these limitations, vibrational imaging, particularly Raman microscopy, offers a promising avenue toward live-cell spatial metabolomics. By probing specific vibrational motions of chemical bonds, Raman scattering provides rich chemical information and enables direct fingerprinting of endogenous metabolites. Through molecular fingerprinting and advanced profiling algorithms, label-free strategies have recently enabled multiplexed analysis in live cells and tissues, with demonstrated correlations to functional transcriptomic readouts termed “RamanOmics”.^18,19^ However, label-free approaches alone are largely limited to static profiling, lacking direct access to metabolic dynamics and offering restricted molecular coverage (e.g., proteins, saturated and unsaturated lipids) due to spectral overlap and background interference.

Recent advances have demonstrated that the integration of small biorthogonal tags (e.g., alkynes, deuterium, nitriles that reside in the ideal cell-silent spectral region) with stimulated Raman scattering (SRS) microscopy provides a powerful strategy to overcome these limitations, enabling minimally perturbative quantitative imaging of metabolic dynamics in live cells with target specificity, subcellular spatial resolution, and high temporal fidelity (**Fig. 1a**).^20,21^ Indeed, metabolic activity phenotyping using several of these bioorthogonal probes has demonstrated much-improved separation and sensitivity compared to purely label-free imaging.^22^ Although the intrinsically narrow Raman linewidths (∼10 cm^-1^) have enabled super-multiplex Raman imaging with up to ∼14 channels through conjugation-based strategies, the resulting probes are typically bulky and therefore unsuitable for labeling small metabolites without perturbing their native functions during imaging and profiling.^23^ Therefore, the accessible Raman metabolic palette remains yet to be fully exploited.

**Figure 1.**
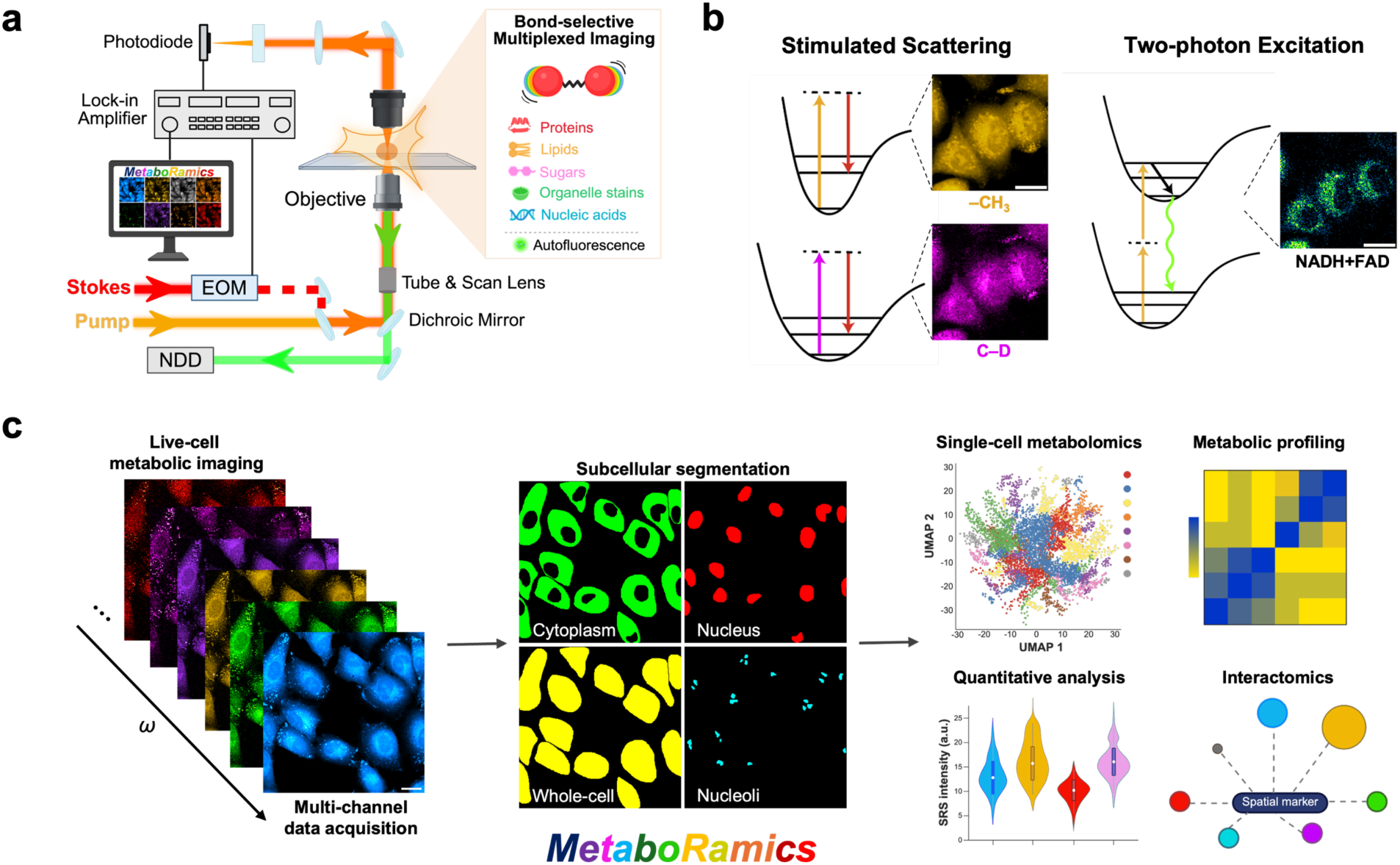
MetaboRamics workflow. **(a)** SRS microscopy setup for bond-selective multiplexed imaging, integrated with two-photon fluorescence detection using the same laser sources. EOM = electro-optic modulator, NDD= non-descanned detector; **(b)** Energy diagram for stimulated Raman scattering (left) and two-photon excitation (right); **(c)** Workflow for spatial metabolomics using MetaboRamics.

Here, we report a robust analytical platform, termed MetaboRamics, for highly multiplexed **metabo**lic imaging by stimulated **Ram**an for spatial metabolom**ics** in live cells (**Fig. 1**). We achieve this through rational probe screening and careful selection to ensure high spectral separation, in combination with isotope editing strategies and robust unmixing algorithms. Together, these advances substantially expand the accessible Raman metabolic palette while maintaining minimal perturbation to native cellular processes. We demonstrate a greatly expanded and minimally invasive metabolic palette for 16-plex MetaboRamics in live cells, integrating nine bioorthogonal channels targeting key metabolic pathways-including sugars, choline, DNA, proteins, lipids, and organelle markers-alongside five label-free channels and two autofluorescence channels from two-photon excited fluorescence (TPEF) microscopy within a unified imaging platform (**Fig. 1a-b**). To fully leverage this multiplexing capability, we further establish a quantitative analysis pipeline with subcellular segmentation, enabling spatially resolved profiling of metabolic states (**Fig. 1c**).

Following systematic optimization and validation, we apply this framework to probe the metabolic basis of epithelial-mesenchymal transition (EMT), a process that remains incompletely understood. Our multi-dimensional analyses reveal system-wide metabolic remodeling where mesenchymal cells display a globally attenuated metabolic state relative to their epithelial counterparts. We further perform subcellular metabolic profiling under diverse metabolic stressors that mimic pathological conditions, including inflammation, serum deprivation, nutrient overload (e.g., high fructose, and saturated fatty acids), and pharmacological treatments to understand their broad effects on metabolic activity. MetaboRamics resolves stressor-specific metabolic rewiring across multiple pathways, revealing both shared and distinct cellular response programs. Collectively, MetaboRamics provides a versatile platform for high-resolution, super-multiplexed imaging of live-cell spatiotemporal metabolomics.

## Results

### 1. Design, Rationale, and Implementation of Live-Cell MetaboRamics Platform

An effective targeted metabolic palette should fulfill the following criteria. First, given the complexity and interconnectedness of metabolic pathways, the palette must encompass a broad range of metabolic species while remaining biocompatible, minimally toxic, cost-effective, and robust for spatial metabolomics. Second, these metabolic analogs should have sufficient uptake and/or incorporation into targeted metabolic pathways with high signal-to-noise ratios (SNR). Third, they should maintain spectral resolvability after incorporation into metabolic pathways. Although the cell-silent region is relatively wide, commonly used vibrational tags (i.e., C–D, alkyne, nitrile) are largely confined to a narrow window (∼2000-2300 cm^-1^). In particular, highly deuterated probes (e.g., d_7_-Glc) exhibit broadened linewidths that fully inundate this region and increase spectral unmixing complexity.

Based on these criteria, we first screened a large set of analogs and isotopologues of glucose, amino acids, lipids and their precursors (**Fig. S1**), yielding a metabolic palette that spans multiple metabolic pathways (**Fig. 2a-b**). Our palette exhibited largely resolvable peaks after metabolic incorporation while retaining high SNR, enabling robust linear unmixing used to deconvolve overlapping channels for downstream quantification (**Fig. 2b**). Specifically, we used 5-ethynyl-2’-deoxyuridine (EdU, 2118 cm^-1^), a thymidine analogue, to map *de novo* synthesis of DNA,^24^ and propargylcholine (PGC, 2142 cm^-1^) for choline metabolism into cellular phospholipids.^25^ We further adopted a monitoring strategy for glucose uptake and metabolism, by synthesizing an isotopologue of an alkyne-tagged glucose analogue, ^13^C_3_-3-OPG (2054 cm^-1^, **Fig. 2a-b**, purple),^26^ to primarily visualize glucose uptake, while in parallel using d_7_-glucose (d_7_-Glc) to track its metabolic incorporation into downstream biomass, including lipids (d_7_-Glc_L_, 2150 cm^-1^) through *de novo* lipogenesis (DNL), and proteins (d_7_-Glc_P_, 2190 cm^-1^).

**Figure 2.**
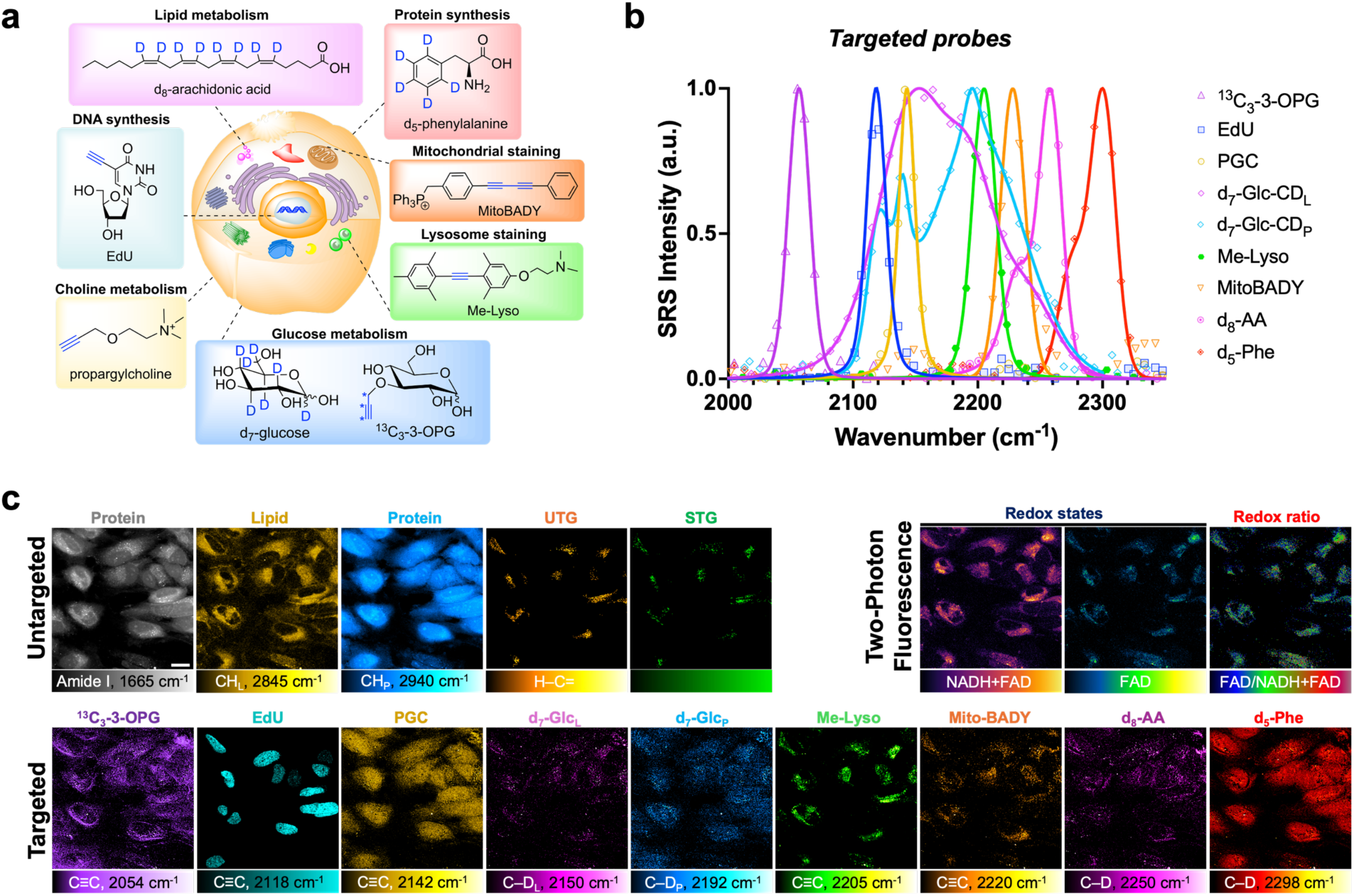
16-plex metabolic imaging using MetaboRamics in live cells. **(a)** Metabolic probe palette encompassing a broad range of labeled metabolites for targeted SRS imaging. **(b)** SRS spectra of incorporated Raman probes from live cells; **(c)** Representative 16-plex metabolic images from live HeLa cells using MetaboRamics. Scale bar: 20 µm.

Extending into the unoccupied cell-silent spectral region, we incorporated key spatial organelle markers, including Me-Lyso (ML, 2205 cm^-1^) to label lysosome^27^ and Mito-BADY (MB, bearing a conjugated diyne, 2220 cm^-1^) as a mitochondria tracker,^28^ that enabled us to assess both organelle dynamics and metabolite-organelle interactomics using our platform. To complement the DNL readout from d_7_-Glc, we selected d_8_-arachidonic acid (d_8_-AA, 2250 cm^-1^) to assess the uptake of polyunsaturated fatty acid (PUFA). We note that d_8_-AA was used at low concentrations (i.e. 2 µM) throughout, as higher concentrations promotes lipid droplet formation, complicating multi-channel analysis due to increased cross-phase modulation (**Fig. S2**). Lastly, we employed d_5_-phenylalanine (d_5_-Phe, C_sp2_–D, 2298 cm^-1^, **Fig. 2b**, red), an essential amino acid positioned outside congested spectral region, to visualize nascent protein synthesis. To further expand the palette into less crowded spectral windows, a potentially viable design is to exploit deuterated terminal alkynes (C≡C–D), which exhibits a peak at ∼1985 cm^-1^.^27^ However, although we synthesized 17-D-ODYA to probe SFA uptake, we observed relatively rapid hydrogen-to-deuterium exchange (HDX), resulting in partial conversion back to protonated alkyne (C≡C–H, 2120 cm^-1^) in cells, rendering it unsuitable for this application (**Fig. S3**).

With these nine-color targeted probes enabling the simultaneous interrogation of metabolic dynamics (**Fig. 2c, bottom**), we next integrated them with established label-free channels, including amide I (C=O stretching of peptide backbone, 1665 cm^-1^), lipids (–CH_2_, 2845 cm^-1^), proteins (–CH_3_, 2940 cm^-1^) and unsaturated lipids (=C–H, 3022 cm^-1^). Here, saturated triglycerides (STG) in lipid droplets (LDs) were quantified using a previously established approach, in which unsaturated triglycerides (UTG) are subtracted from the total lipid pool (CH_2_) to yield STGs (**Fig. S4**), hence providing five label-free channels (**Fig. 2c, top left**).^29^ In parallel, label-free TPEF microscopy was employed to optically read out the cellular redox state by capturing endogenous autofluorescence from the redox coenzymes NADH and FAD, using the SRS pump laser for excitation (NADH is excited only at 760 nm; whereas FAD is excited at both 760 nm and 880 nm) and with non-descanned detectors (NDD) employed for signal collection (**Fig. 2c, top right**). The resulting NADH+FAD (reduced + oxidized) to FAD (oxidized) ratio serves as a functional readout of cellular metabolic and mitochondrial activity, as both coenzymes are closely linked to mitochondrial oxidative metabolism.

With this framework, MetaboRamics enables, for the first time, rapid, high-throughput 16-plex imaging in live HeLa cells (**Fig. 2c, Fig. S5**). Each high-resolution image set was acquired within ∼25 minutes, primarily limited by pump-laser tuning, while inducing negligible photodamage. Control comparison between spectrally unmixed 16-plex images with corresponding single-channel SRS images, in which each probe was administered individually (**Fig. S6**), confirm faithful preservation of spatial features, demonstrating minimal probe-associated interference and toxicity.

### 2. MetaboRamics Reveals Metabolic Reprogramming in EMT

With our ability to perform 16-plex metabolic imaging, we next applied MetaboRamics to investigate the metabolic changes underlying EMT that remain incompletely characterized. EMT is a process by which epithelial (epi) cells become de-differentiated into mesenchymal (mes) cells, losing cell-to-cell adhesion and polarity while becoming more migratory in nature.^30^ Though EMT has been linked with essential processes such as embryonic development and wound healing, it has also been reported to progress pathologic responses like cancer metastasis.^31^ However, most studies linking EMT with altered metabolism rely on bulk measurements or invasive methods that do not capture metabolic intracellular heterogeneity in living systems.^32^ Recent studies using optical imaging of live cells have provided insights into metabolic changes associated with EMT.^33,34^ However, a more holistic characterization of the metabolic landscape of EMT *in situ* still remains lacking.

To model EMT, we first induced EMT in A549 epi cells via a commercial cocktail of proteins and antibodies. We successfully performed high-quality, 16-color MetaboRamics imaging on both epithelial and mesenchymal phenotypes (**Fig. 3a**) and captured clear morphological transition in the label-free protein imaging (CH_P_, 2940 cm^-1^), where epi cells exhibited a cobble-stone morphology, while mes cells became elongated with increased fenestrations (**Fig. 3a**). To quantitatively visualize any present metabolic changes, we next constructed a population-averaged heatmap, plotting the log_2_fold-change (log_2_FC) at the whole cell and subcellular level (segmented by cytoplasm, nuclei, and nucleoli) (**Fig. 3b**), standardized against A549 epi (control) cells.^34^ We found that EMT showed a global reduction by ∼30% across all probed metabolic pathways in, consistent with our prior findings on TCA-linked metabolism.^34^ (**Fig. 3b**).

**Figure 3.**
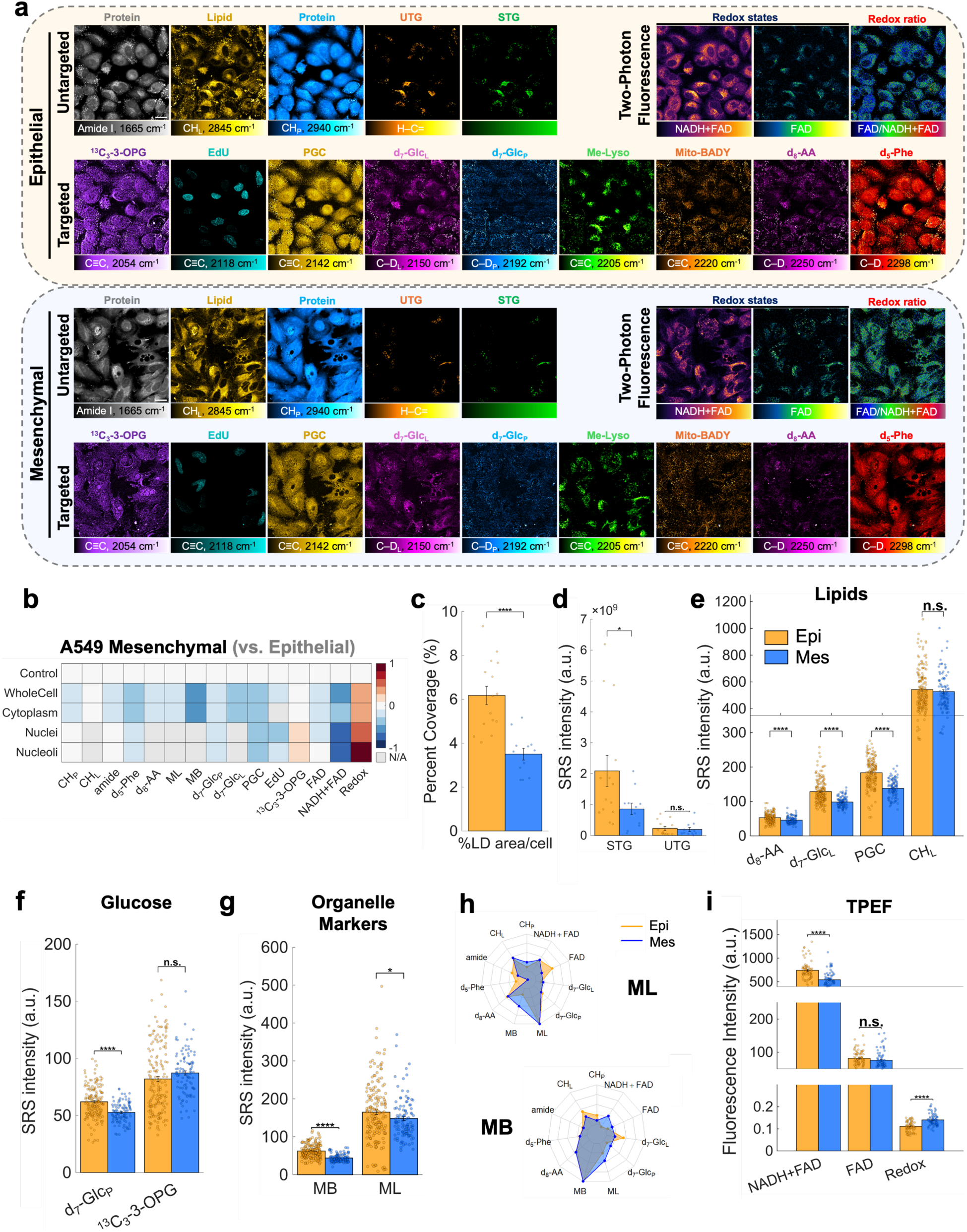
MetaboRamics Reveals Metabolic Reprogramming in EMT. **(a)** Representative 16-plex metabolic images using MetaboRamics in live A549 epithelial (top) and mesenchymal (bottom) cells. Scale bar: 20 µm; **(b)** Heatmap depicting log_2_ fold changes between epithelial and mesenchymal cells profiled by MetaboRamics at the subcellular level (whole-cell, cytoplasm, nuclei and nucleoli). The control comprises of epithelial cells; **(c)** Percent area coverage of LDs per field-of-view (FOV) in EMT; **(d)** Quantification of SRS signal (intensity x area) per FOV of saturated triglycerides (STG) and unsaturated triglycerides (UTG); **(e)** Quantification of SRS intensity per cell for CH_L_ (endogenous total lipids), d_8_-AA (lipid uptake), d_7_-Glc_L_ (*de novo* lipogenesis) and PGC (choline metabolism) for EMT; **(f)** Subcellular quantification of SRS intensity per cell for d_7_-Glc_P_ (protein synthesis) across the whole-cell, cytoplasm, nuclei and nucleoli; **(g)** Quantification of SRS signal per cell for organelle markers: MB and ML; **(h)** Radial plots for MB (top) and ML (bottom) depicting pairwise interaction networks across channels; **(i)** Quantification of TPEF signal per cell from NADH + FAD, FAD, and redox ratio. Data represented as mean ± SEM from at least three independent experiments. Statistical analysis was conducted using Welch’s two-paired t-test, *p<0.05, **p<0.01, ****p<0.0001 and n.s.= no statistical difference.

Surprisingly, label-free lipid channel images (CH_L_; **Fig. 3a,c** and **Fig. S7**) revealed a marked decrease (∼30%) in the number of LDs in the mes phenotype. This trend contrasts with previous observations in MCF7 cells, where mes cells exhibited increased LD abundance relative to their epi counterparts,^33^ highlighting substantial cell-type-dependent differences in metabolic reprogramming during EMT. Interestingly, composition analysis showed that STGs in LDs were significantly reduced (∼2 fold), whereas UTGs remain relatively constant (**Fig. 3d**). Our further analysis revealed decreased DNL (i.e. lower d_7_-Glc_L_ signals) and PUFA uptake and metabolism (i.e. lower d_8_-AA signal) in mes cells (**Fig. 3b,e**). In addition, choline metabolism for phospholipid synthesis was also diminished (**Fig. 3e**, PGC), collectively suggesting a global downregulation of lipid biosynthetic activity. Despite this reduction in DNL and lipid uptake in mes cells, the unchanged intensity from endogenous lipid channel including LD’s (i.e. CH_L_, **Fig. 3b,e**) indicates that the total lipid biomass is largely maintained. This apparent balance could likely be attributed to our observed enhanced mobilization of lipids from LDs, suggesting that mes cells sustain lipid homeostasis through increased lipid recycling and decreased degradation despite a globally attenuated metabolic state.

In addition to lipid remodeling, MetaboRamics also revealed interesting differences in glucose dynamics in EMT. Our data shows that glucose metabolism for both lipid and protein synthesis (d_7_-Glc_L_ and d_7_-Glc_P_, respectively) is reduced in mes cells (**Fig. 3b, e, f**), whereas glucose uptake (^13^C_3_-3-OPG) remained relatively unchanged (**Fig. 3b, f**). These results indicate that, despite comparable glucose uptake between epi and mes cells, glucose metabolism via glycolysis in mes cells is less effectively channeled into biosynthetic pathways. As a control, we validated the glucose uptake trend using single-color SRS imaging for EMT with ^13^C_3_-3-OPG alone, which yielded consistent results (**Fig. S8**), again confirming minimal cross-channel interference among probes.

Organelle markers further highlighted state-dependent differences during EMT (**Fig. 3b,g**). We observed a decrease in MB signal in mes cells which may reflect reduced mitochondrial abundance and/or organization (e.g., fragmentation vs. networked structure),^34^ consistent with our prior observations that EMT attenuates mitochondrial TCA-cycle metabolism.^34^ Similarly, a modest attenuation in ML signal was detected in mes cells, suggesting potential differences in lysosomal abundance, distribution, and/or size (**Fig. 3b,g**). Our spatial interactome analysis further uncovered EMT-dependent organelle coupling (**Fig. 3h**). In epi cells, lysosomes (i.e. ML) showed stronger association with FAD and protein synthesis markers (d_5_-Phe), whereas mitochondria (i.e. MB) preferentially coupled to lipid-related signals (CH_L_ and d_7_-Glc_L_) (**Fig. 3h**, orange). Consistently, quantification of TPEF signals revealed minimal changes of FAD intensity between epi and mes cells, but a substantial decline in the NADH + FAD mixed channel, indicating reduced NADH levels in mes cells and, consequently a more oxidized mitochondria state^35^ in EMT (**Fig. 3i**). Taken together, our multi-channel MetaboRamics analysis reveals that mes cells exhibit reduced glucose incorporation into biomass, altered mitochondrial activity and decreased NADH levels compared with epithelial cells, reflecting a global metabolically attenuated state.

### 3. High-throughput Metabolic Phenotyping of Stress-induced Cellular Responses

Going beyond EMT, to systematically probe cellular metabolic responses under disease-relevant conditions, we curated a set of nine metabolic stressors spanning nutrient excess, serum deprivation, inflammation, and pharmacological perturbations. To model nutrient overload, we subjected A549 cells to high fructose (FRUC) which contributes to the pathogenesis of cardiometabolic disease such as diabetes, obesity, and metabolic fatty liver disease.^36^ Additionally, we chose palmitic acid (PA, as a prototypical SFA) which causes lipotoxicity and endoplasmic reticulum stress.^37^ Moreover, to mitigate growth-factor-dependent metabolic activity, we used FBS-free media for serum deprivation. Furthermore, cells were subjected to TGF-β and IL1-β to mimic inflammation (INF) and trigger EMT.^38^ Since inflammation often accompanies the ramifications caused by high fructose in other cell types,^39^ we also assessed the effect of the combination of both stimuli (FRUC+INF). Lastly, we pharmacologically modulated protein levels by inhibiting synthesis with cycloheximide (CHX),^40^ blocking degradation with MG132^38^ or bortezomib (BTZ),^41^ and promoting it using CHIR-99021 (CHIR) via GSK-3 inhibition.^42^

As an initial demonstration of our 16-plex metabolic imaging framework, we first implemented a 4-plex imaging strategy (PGC, d_7_-Glc, d_8_-AA and d_5_-Phe) to capture baseline metabolic phenotypes before expanding to higher multiplexing. We constructed a heatmap for all 9 metabolic stressors (**Fig. 4a**). We found that, from nutrient stress, cells subjected to high fructose (i.e. FRUC) displayed an attenuation in PUFA (i.e. d_8_-AA) uptake, likely indicating a lower metabolic demand for fatty acid uptake due to increased DNL from excess fructose. Our data also uncovered that high PA caused a downregulation of lipid-associated metabolism from d_7_-Glc and PGC but stimulated an upregulation of PUFA (i.e. d_8_-AA) uptake (**Fig. 4a**). This observation is in agreement with previous reports^43,44^ that unsaturated fatty acids (UFA) may rescue cells from SFA-induced lipotoxicity and ER stress by sequestration of excess SFAs into TGs within LDs. In contrast, serum deprivation (FBS-free) caused a downregulation in protein synthesis (d_5_-Phe) and PUFA uptake (d_8_-AA) but an increase in biomass synthesis from d_7_-Glc and PGC (**Fig. 4a-c**).

**Figure 4.**
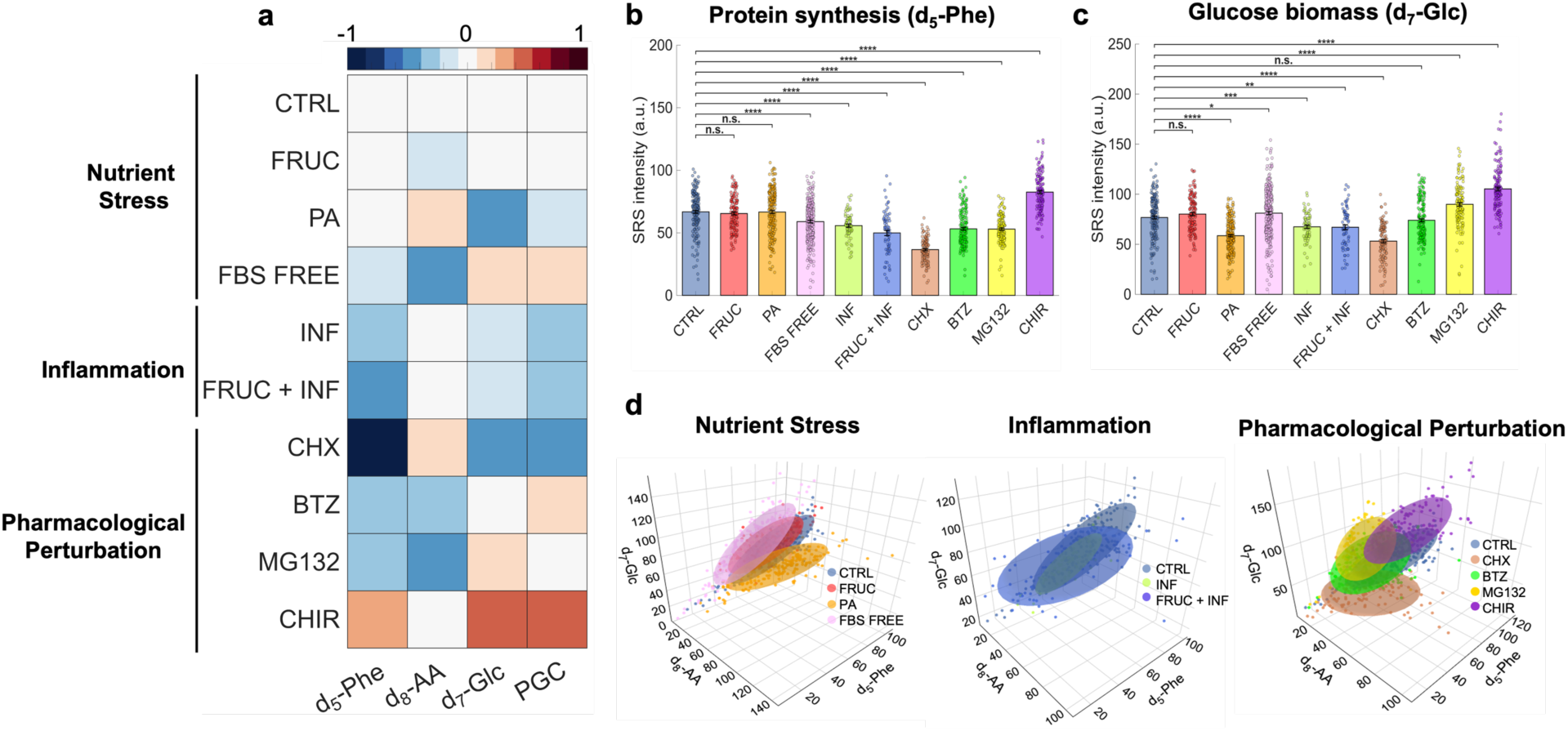
High-content Metabolic Phenotyping of Stress-induced Cellular Responses. **(a)** Heatmap showing log_2_ fold changes from 4-plex SRS imaging (d_5_-Phe, d_8_-AA, d_7_-Glc, PGC) comparing live A549 control cells with cells exposed to nine metabolic stressors (FRUC, INF, FRUC+INF, FBS-free, PA, CHX, BTZ, MG132, or CHIR). Representative SRS images are shown in **Fig. S9**; **(b-c)** Quantification of C–D intensity per cell of **(b)** d_5_-Phe incorporation for protein synthesis and **(c)** d_7_-Glc for glucose-derived biomass in stressor-treated cells; **(d)** 3D scatter plots representing SRS intensities per cell from d_7_-Glc, d_8_-AA, and d_5_-Phe in control and cells treated with different metabolic stressors. Data represented as mean ± SEM from at least three independent experiments. Statistical analysis was conducted using Welch’s two-paired t-test, *p<0.05, **p<0.01, ***p<0.001, ****p<0.0001 and n.s.= no statistical difference.

For inflammatory stimulus (INF), we observed that similar to EMT, this condition caused a downregulation in d_7_-Glc, PGC and d_5_-Phe metabolism but not PUFA (i.e. d_8_-AA) uptake (**Fig. 4a-c**). Interestingly, we found that the combination of FRUC+INF downregulated metabolism of the 4-channels (similar to INF) more profoundly than high fructose (FRUC) alone (**Fig. 4a-c**), suggesting a compounding effect. Furthermore, we found that the additional label-free –CH_3_ channel revealed that the INF condition induced pronounced cellular elongation largely resembling the EMT condition. However, it also differs from EMT through strong lipid accumulation in the cytoplasm, suggesting an overall distinct cellular state from the mesenchymal phenotype altogether (**Fig. S9**).

Pharmacological perturbations led to the most extreme metabolic alterations in A549 cells. We observed a significant decrease in C–D signals from d_5_-Phe (2298 cm^-1^) incorporation upon treatment with protein synthesis inhibitors (CHX [∼45%], MG132 [∼21%], BTZ [∼20%]) (**Fig. 4b**, CHX/MG132/BTZ vs CTRL). As a comparison, CHIR treatment displayed broad upregulation, including the largest increase (∼25%) in d_5_-Phe derived signals across all our series of metabolic stressors (**Fig. 4b** CHIR vs CTRL). We note that these drug perturbations indeed have widespread and unexpected metabolic influences. For example, while CHX, MG132, and BTZ are all protein synthesis inhibitors, MG132 upregulates glucose metabolism (**Fig. 4c**, MG132 vs CTRL), whereas CHX strongly downregulates glucose metabolism (**Fig. 4c**, CHX vs CTRL ) and BTZ had minimal effect on glucose metabolism (**Fig. 4c**, BTZ vs CTRL ), highlighting their fundamentally varied mechanisms of action (MoA) on a larger biological network, which are worth comprehensively investigating when evaluating drug effects, underscoring the value of our platform. Taking advantage of the high dimensionality of our single-cell multiplexed measurements, we then mapped multiple metabolites into a three-dimensional chemical space (**Fig. 4d**), visualizing clustering due to the differential responses of d_7_-Glc, d_8_-AA, and d_5_-Phe to each conditions; consistent with previous reports, such clustering again highlight different MoA of the drugs.^45^ Combining the high biocompatibility of small vibrational tags with sensitive SRS microscopy, 4-plex metabolic imaging thus already robustly allows high-throughput screening, enabling systematic characterization of varied drug responses and interactions.

### 4. Optical Phenotyping of Cellular Stress Responses Using MetaboRamics

Seeking to further disentangle the complex and contrasting metabolic profiles obtained from treatment with CHIR, CHX, and INF, we applied our high-content, 16-plex MetaboRamics platform to thoroughly investigate their metabolic responses in live A549 cells at both single-cell and subcellular levels. We constructed a log_2_FC heatmap at both whole-cell and subcellular levels, organized by subcellular compartments (i.e. cytoplasm, nuclei, and nucleoli) and treatment conditions (i.e. CHIR, CHX, and INF) (**Fig. 5a**). Our whole-cell super-multiplexed data exhibited excellent agreement with the 4-plex data (**Fig. S10**), highlighting the robustness and minimal cellular perturbation of our platform. For example, upon treatment with CHIR, we again observe a global upregulation among the metabolic probes reporting *de novo* synthesis, and uptake (**Fig. 5a-c**, CHIR, orange). Interestingly, our multiplexed imaging, however, reveals a significant downregulation of both ML and MB (**Fig. 5d**, left; CHIR, orange), suggesting complex perturbation mechanisms to mitochondrial and lysosomal activities. As another example, CHX and INF treatment resulted in similar metabolic profiles across metabolic labels, with general attenuation across the targeted channels, but CHX upregulated both ML and MB whereas INF did not affect MB signal (**Fig. 5d**, left). These findings further highlight the need for parallel global organelle-specific analysis during metabolic stress.

**Figure 5.**
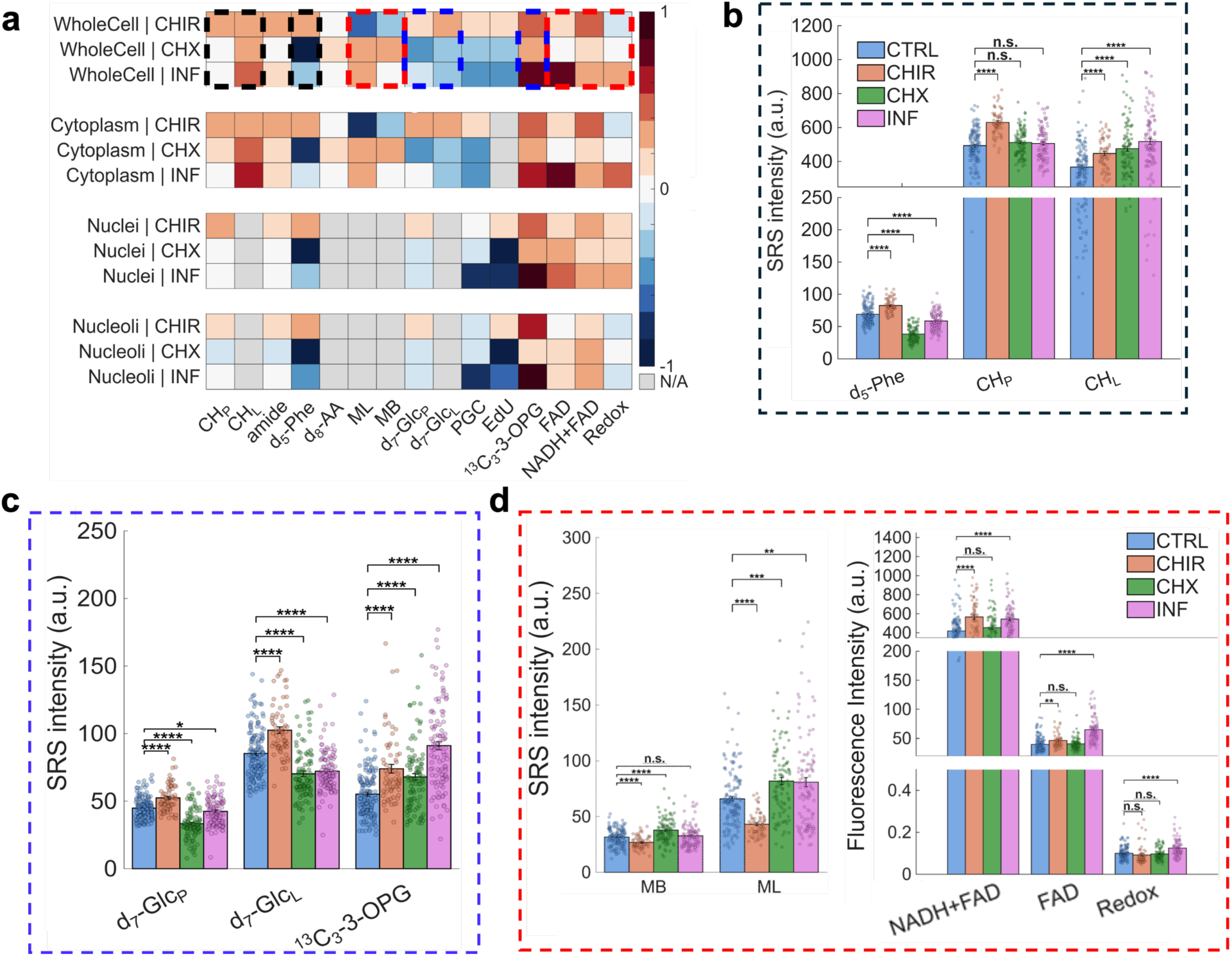
Optical Phenotyping of Cellular Stress Responses Using MetaboRamics. **(a)** Heatmap from super-multiplexed metabolic imaging reflecting log_2_ fold changes between the A549 control and cells treated with CHIR, CHX, or INF at the subcellular level (whole-cell, cytoplasm, nuclei and nucleoli); **(b)** Quantification of SRS intensities per cell for (left to right) d_5_-Phe, CH_P_ and CH_L_ across different conditions: CTRL (blue), CHIR (orange), CHX (green) and INF (magenta); **(c)** Quantification of SRS intensities per cell for glucose metabolism (d_7_-Glc_p_ and d_7_-Glc_L_) and glucose uptake (^13^C_3_-3-OPG). **(d)** (Left) Quantification of SRS intensities per cell for organelle markers (MB and ML); (Right) TPEF intensities per cell for NADH+FAD, FAD, and redox ratio. Data represented as mean ± SEM from at least three independent experiments. Statistical analysis was conducted using Welch’s two-paired t-test, *p<0.05, **p<0.01, ***p<0.001, ****p<0.0001 and n.s.= no statistical difference.

Indeed, metabolic labeling enables direct interrogation of dynamic metabolic activity whereas label-free analysis provides complementary insights into steady-state cellular composition. For instance, using d_5_-Phe to probe nascent protein synthesis, we observed a higher inhibitory effect with CHX than with INF. In contrast, the label-free CH_p_ channel showed no significant change in overall protein content across treatments, suggesting a slower degradation upon the drug treatments (**Fig. 5b**; CRTL (blue) vs CHX (green) and INF (magenta)). Interestingly, while CHIR upregulated d_5_-Phe metabolic signals, the increase in the label-free CH_P_ channel was even larger (20% vs 27%), suggesting that, in addition to enhanced protein synthesis, protein degradation may also be attenuated (**Fig. 5b**; CTRL (blue) vs CHIR (orange)). At the subcellular level, we further found protein modulation resulted in spatially heterogenous metabolic changes. For instance, we found CHX uniformly degrades protein synthesis across all cellular compartments (whole-cell, cytoplasm, nuclei and nucleoli) but INF causes pronounced changes in its nucleoli (i.e., higher than the cytoplasm) whereas CHIR had the highest increase in protein upregulation in its nuclear compartments (i.e., both nuclei and nucleoli) (**Fig. 5a** and **S11**).

We next examined our full glucose palette (d_7_-Glc_P_, d_7_-Glc_L_, ^13^C_3_-3-OPG), towards deeper insights into glucose dynamics. Consistent with our 4-plex experiments (**Fig. 4a**), CHIR upregulates d_7_-Glc_P_ and d_7_-Glc_L_, while CHX and INF downregulate d_7_-Glc_P_ and d_7_-Glc_L_ (**Fig. 5c**). Surprisingly, ^13^C_3_-3-OPG signal increased for all three metabolic stressors, indicating a relative decoupling of glucose uptake (increased ^13^C_3_-3-OPG) and downstream metabolic incorporation (decreased d_7_-Glc_P_ and d_7_-Glc_L_) for CHX and INF (**Fig. 5c**). These results further contrast with our EMT measurements (**Fig. 3b**), where glucose uptake (^13^C_3_-3-OPG signal) remained unchanged despite reduced glucose metabolism (decreased d_7_-Glc_P_ and d_7_-Glc_L_). These observations suggest differences in uptake dynamics and metabolism unique to these perturbations.

Complementary TPEF imaging revealed increases in absolute intensity for both FAD and NADH+FAD in CHIR- and INF-treated cells (**Fig. 5d**, right). Analysis of the redox ratios (FAD/(NADH+FAD)) revealed that CHX (green) and CHIR (orange) induced minimal changes, whereas INF (magenta) shifted cells toward a more oxidized state, as reflected by an increased redox ratio (**Fig. 5d**, right). This shift is consistent with altered mitochondrial balance, potentially reflecting enhanced oxidative metabolism or changes in electron transport dynamics under INF treatment. These trends partially align with glucose-derived metabolic readouts (**Fig. 5c**), where INF (magenta) treatment is associated with decreased lipid and protein anabolism (d_7_-Glc_L_ and d_7_-Glc_P_). However, this relationship diverges under CHIR, which exhibits enhanced anabolic activity despite minimal changes in redox ratio, suggesting a partial decoupling between mitochondrial redox state and biosynthetic output. Together, our results highlight the importance of multiplexed metabolic imaging to capture distinct and partially uncoupled aspects of cellular metabolism.

## Discussion

Metabolism is a highly spatially organized series of chemical reactions, yet most live-cell imaging methods only capture a narrow portion of metabolic activity at a time. To maximize the information rendered from optical imaging methods for spatial metabolomics, it is necessary to simultaneously probe several different metabolic pathways and examine how they interact with each other. Here, we demonstrate MetaboRamics as a super-multiplexed SRS imaging platform that enables simultaneous, spatially resolved mapping of metabolic activity in live cells. In addition to mapping endogenous biomolecules such as proteins and lipids (total lipids, UTG, and STG), our method captures a broad range of metabolic pathways: glucose uptake, glucose metabolism that leads to nascent synthesis of biomass including lipids and proteins, *de novo* DNA synthesis, choline metabolism, fatty acid uptake, and *de novo* protein synthesis. With the use of spatial markers for organelles such as lysosomes and mitochondria, our analytical workflow also enables interactomics to understand the interplay between cellular organelles and metabolites. Finally, the highly complementary TPEF provides an optical reporter for the redox states of cells, achieving 16-plex imaging in live HeLa with MetaboRamics.

We next applied our 16-plex platform to investigate the metabolic landscape of EMT in live A549 cells, which remains incompletely characterized. Our multi-dimensional data indicated a global attenuation of metabolic activity in several metabolic pathways in mes cells compared to their epithelial counterparts. For instance, through a decrease in DNL and PUFA uptake, we found a reduction in lipid turnover in mes cells. Additionally, using two different glucose reporters-we revealed that despite having similar glucose uptake, mes cells have lowered glucose-derived biosynthetic activity. By integrating complementary TPEF, we further demonstrated that mes cells accumulate NADH due to reduced NADH oxidation rather than increased glucose flux which is consistent with a change in redox balance, and global metabolic attenuation. Our method highlights the metabolic flexibility displayed by mesenchymal cells to become quiescent which in many cases has been attributed to the drive chemoresistance to therapeutic agents.^46^

We next performed high-content metabolic phenotyping to characterize cellular responses under disease-relevant conditions. MetaboRamics revealed consistent, pathway-specific metabolic adaptations across diverse metabolic stressors. For example, similar to EMT, inflammatory stimuli led to a global downregulation of metabolism in A549 epithelial cells, with select exceptions, including increased ML, glucose uptake and lipid metabolism. Similarly, CHX selectively downregulated *de novo* protein synthesis in cells but increased PUFA uptake. Therefore, these results demonstrate the ability of our platform to simultaneously capture integrated metabolic responses rather than isolated pathway changes.

We also note a few limitations. Although our current probe panel covers a broad range of metabolic pathways, it does not fully encompass the full diversity of cellular metabolism, largely constrained by overlapping Raman spectra and leading to challenging unmixing. Additionally, our imaging speed for 16 channels is limited by laser tuning requirements of our cavity-based picosecond laser, restricting the total time taken for multiplexing. The use of rapidly tunable lasers could enable faster image and spectral acquisition that would also facilitate the use of more sophisticated unmixing tools such as LASSO.^47,48^ Furthermore, the integration of deep learning algorithms can also assist in faster image acquisition (or image prediction) by drastically increasing the signal-to-noise ratio of SRS images, requiring lower laser power and reducing photodamage as more channels are acquired.^49^ Lastly, despite the use of Raman tags, the sensitivity of SRS is still limited in the low µM to low mM regime.^50^ This constraint poses challenges for many metabolic species that are present at substantially lower concentrations in cells (in ∼nM ranges). Future efforts to engineer metabolic probes that leverage epr-SRS^51^ to improve detection limits would greatly expand the ability to visualize sub-micromolar species.

To perform quantitative spatial metabolomics in absolute concentrations, future work should combine MetaboRamics with newly developed frameworks such as MATRIX-SRS^34^ that can provide absolute concentrations for deuterium-based biomass. Additionally, one can envision a multi-omics platform that leverages the strong complementarity of MetaboRamics with other modalities, such as spatial transcriptomics (e.g., Visium)^52^ or single-cell RNA sequencing^18^, on the same sample to link transcriptional changes with metabolic states and gain deeper systems-level understanding of cellular function. Similarly, coupling MetaboRamics with MSI, such as MALDI,^50^ could enable deeper metabolome coverage. Lastly, beyond fundamental studies of metabolism, MetaboRamics is also well suited for applications in drug discovery: to probe multiple metabolic outcomes, identify off-target effects, and characterize the MoA of drug candidates. Furthermore, our methodology has the potential to be applied as an optical pooled screening^53^ method for metabolic phenotyping of large-scale pooled perturbation (e.g., CRISPR) screens in mammalian cells.

Overall, our work demonstrates the ability of MetaboRamics to resolve metabolic complexity and spatial heterogeneity in live cells, providing a versatile framework to study metabolic states and rewiring across biological contexts.

## Materials & Methods

### Cell Culture and Metabolic Labeling

HeLa cells (ATCC, CCL-2), and A549 cells (gift from the Elowitz lab, Caltech) were each cultured in DMEM (Gibco, Cat. #11965092) supplemented with 1x antibiotic-antimycotic (Gibco, Cat. #15240062) and 10% FBS (Corning, Cat. #35-015-CV) at 37 °C with 5% CO_2_. For EMT induction, A549 cells were seeded and cultured in DMEM supplemented with 1x StemXVivo EMT-inducing supplement (R&D Systems, Cat. #CCM017) for 5 days, with media changes every 2-3 days.

Metabolic labels used for MetaboRamics included 1 mM propargylcholine bromide (MedChemExpress, Cat. #HY-129084), 25 mM 1,2,3,4,5,6,6-d_7_-D-glucose (Cambridge Isotope Laboratories, Cat. #DLM-2062), 1 mM d_5_-L-phenylalanine (Cambridge Isotope Laboratories, Cat. #DLM-1258-5), and 25 mM ^13^C_3_-3-OPG (synthesized according to previously reported method^26^), which were added to cells in custom DMEM with no unlabeled counterparts for 48 hours. Following 24 hours, 200 µM 5-Ethynyl-2’-deoxyuridine (MedChemExpress, Cat. #HY-118411) and 2 µM d_8_-arachidonic acid (MedChemExpress, Cat. #HY-109590S) were added for the remaining 24 hours. The Me-Lyso stock solution (in DMSO) was photo-uncaged by irradiation with a 405 nm laser for 10 min prior to incubation with cells. Finally, 400 nM MitoBADY and 7.5 µM Me-Lyso (see Supplementary Information for synthetic protocols) were added to the cells for 30 minutes before multi-channel SRS imaging. These concentrations and incubation times were selected following systematic optimization across labels to ensure high imaging signal-to-noise ratios while maintaining high cellular viability.

Metabolic stressors included 25 mM D-fructose (FRUC; Thermo Fisher, Cat. #A17718.0I), 10 ng/mL IL-β1 (R&D systems, Cat. #201-LB-005) and 10 ng/mL TGF-β (R&D systems, Cat. #7754-BH-005) (INF), 25 mM D-fructose + 10 ng/mL IL-β1 and 10 ng/mL TGF-β (FRUC + INF), 100 µM palmitic acid (PA; Sigma-Aldrich, Cat. #506345-25GM), 3.5 µM CHIR99021 (CHIR; MedChemExpress, Cat. #HY-10182), 5 µM MG132 (MedChemExpress, Cat. #HY-13259), 1.7 µM Cyclohexamide (CHX; Sigma Aldrich, Cat. #239763-1GM), 10 nM Bortezomib (BTZ; MedChemExpress, Cat. #HY-10227) and DMEM containing no FBS (FBS-free). For FRUC, INF, and their combinatorial treatment (FRUC + INF), A549 cells were first exposed to these stressors alone for 48 hr before the concomitant addition of metabolic labels for 24 hr. All other stressors were added concomitantly with the metabolic labels and incubated for 24 hr prior to SRS imaging.

### SRS Microscopy

A tunable pump beam (700-990 nm, 2 ps, 80 MHz) and Stokes beam (1032.0 nm, 2 ps, 80 MHz, modulated to 20 MHz by an Electro-Optic modulator, EOM) from a picoEmerald S laser system (Applied Physics & Electronics) are used as the light source for SRS Microscopy. The spatially and temporally overlapped beams are expanded and introduced into an inverted multiphoton laser scanning microscope (FV3000, Olympus). Light is focused on the sample with a 25x water immersion objective (XLPLN25XWMP, 1.05 N.A., Olympus) and transmitted light is collected with an oil immersion condenser lens (1.40 N.A., Olympus). The Stokes beam is filtered out (893/209 BrightLine, 25 mm, Semrock) and the remaining pump beam is collected with a large area photodiode (S3590-09, Hamamatsu). A 64 V reverse-bias DC voltage was applied to the photodiode to increase the saturation threshold and reduce the response time. The output current was terminated by a 50 Ω terminator and prefiltered by a 19.2-23.6 MHz band-pass filter (BBP-21.4+, Mini-Circuits) to reduce laser and scanning noise. The SRS signal within the pump beam is demodulated via lock-in amplification (SR844, Stanford Research Instruments) at the reference modulation frequency, and the in-phase X input was fed back to the IO interface box (FV30-ANALOG). Images were captured using Olympus FluoView 3000 software.

Images were acquired with an 80 µs pixel dwell time to achieve an image acquisition speed of 8.52 s per frame for a 320×320-pixel field of view (FOV) at a 0.497 µm/pixel resolution. The pump beam wavelength was set to 787.0 nm for the unsaturated triglyceride, 791.8 nm for the protein (–CH_3_ channel, 2940 cm^-1^), 797.8 nm (–CH_2_ channel, 2845 cm^-1^), 834.2 nm (C–D channel for d_5_-phenylalanine, 2298 cm^-1^), 837.5 nm (C–D channel for d_8_-arachidonic acid, 2250 cm^-1^), 839.6 nm (C≡C channel for Mito-BADY, 2220 cm^-1^), 840.7 nm (C≡C channel for Me-Lyso, 2205 cm^-1^), 841.6 nm (C–D protein channel for d_7_-glucose 2192 cm^-1^), 844.6 nm (C–D lipid channel for d_7_-Glucose, 2150 cm^-1^), 845.2 nm (C≡C channel for propargylcholine, 2142 cm^-1^), 846.6 nm (C≡C channel for EdU, 2118 cm^-1^), 851.5 nm (C≡C channel for ^13^C_3_-OPG, 2054 cm^-1^), 855.4 nm (off-resonance background image, 2000 cm^-1^), 880.6 nm (amide I band, 1665 cm^-1^) All images were processed and color-coded using Fiji (ImageJ 2.16.0; ImageJ 1.54p)^54^ software. For hSRS of solutions, the wavelength of the pump laser was tuned from 834.0 nm to 855.0 nm with a step size of 0.5 nm.

For C–D quantification, on and corresponding off-resonance images were preprocessed with the Fiji ‘nonuniform background removal’ plugin before subtracting them, resulting in the C–D (on-off) images that are used for linear unmixing. For label-free quantification, only the Fiji ‘nonuniform background removal’ plugin was used. For cellular and subcellular quantification, the protein (CH_3_) image was used to segment each cell and its compartments utilizing the ‘IdentifyObjectsManually’ module in Cellprofiler 4.2.8, converted to a mask using the ‘ConvertObjectsToImage’ module and saved with the ‘SaveImages’ module. The mask was then fed into MATLAB (R2024b, MathWorks) for identification with the “bwlabel” function and used as a map for quantification for all mask objects. Results are then exported to MS Excel for analysis. Images and graphs are visualized using Fiji, MATLAB, and RStudio.

### Linear Combination Algorithm for Spectral Unmixing

While single-frequency SRS imaging maps the distribution of individual Raman peaks with high chemical specificity, many vibrational modes overlap.^55^ To deconvolute discrete molecular signatures from overlapping peaks, we employed a linear combination algorithm. All image sets were unmixed using MATLAB (Mathworks, USA), with image segmentation and processing performed in Fiji (ImageJ 2.16.0; ImageJ 1.54p). hSRS spectra for individual probes were extracted from hSRS stacks and used to generate the coefficients for the unmixing matrix.

Single-color hSRS stacks for ^13^C_3_-3-OPG (50mM, 2 day incubation), EdU (200 µM, 1 day), PGC, Me-Lyso (7.5 µM, 30 minutes), MitoBADY (400 nM, 30 minutes), and d_5_-Phe (400 µM, 3 days) were acquired in live HeLa cells incubated with the respective probes, whereas hSRS stacks for d_8_-AA (50 µM, 1 day) were acquired in fixed HeLa cells to mitigate LD drift. After processing hyperspectral images (See **SRS Microscopy**), regions of interest (ROIs) with high SNR were segmented and averaged at each frequency to construct an SRS spectrum for each probe. After treatment with d_7_-glucose (25mM, 3 days), hSRS images of d_7_-glucose-derived lipid (CD_L_) were acquired in live A549 cells. For d_7_-glucose-derived protein (CD_P_) measurements, fixed HeLa cells were washed with 0.5% Triton X-100 (Sigma, T8787) in phosphate-buffered saline solution for 10 minutes at 4°C. LDs and nucleoli were segmented and averaged at each frequency to construct spectra for CD_L_ and CD_P_, respectively. hSRS spectra were baseline-corrected as needed and fit with Voigt line shapes in Fityk (v1.3.1),^56^ followed by normalization to their maximum intensity. A distinct unmixing matrix was prepared for each image set, as the exact pump wavelength varies during image acquisition. SRS intensities at each Raman shift were extracted from the spectra and used as coefficients for the unmixing matrix. Each image set is then unmixed with a distinct matrix, either 9 x 9 or 4 x 4 depending on the sample treatment (**Fig. S5**).

To ensure the fidelity of unmixed data, any pixels having a negative contribution of ^13^C_3_-3-OPG, PGC, CD_L_, d_8_-AA, or d_5_-Phe were discarded from all channels prior to analysis. The spatially distinguishable probes–EdU, MB, and ML– were exempted to preserve a data set large enough for analysis, representing a meaningful limitation to the methodology.

For high-frequency C–H stretching region, the lipid channel (CH_L_) was generated using a linear combination of the CH_2_ and CH_3_ images = 5·[CH_2_] − 0.4·[CH_3_] whereas the protein channel (CH_P_) was generated using a linear combination of [CH_3_] − [CH_2_] from the same set of images.^44^

Since the EdU signal is confined to the nucleus, unmixed EdU images were multiplied by nuclear masks to suppress contributions from non-nuclear regions (e.g., cross-phase modulation from LDs) and mitigate background noise.

STG images were calculated according to the workflow described in **Fig. S4**.

### Two-Photon Fluorescence (TPEF) Microscopy

The same picoEmerald used for SRS microscopy was also used to perform TPEF microscopy with the addition of a home-built light-shielding box. Label-free excitation of FAD and NADH was performed at 760 nm, and the separate excitation of FAD was performed at 880 nm. The backscattered light is filtered with a 690 nm short-pass filter and epi-autofluorescence is collected through a non-descanned detector (NDD). TPEF images were collected at the same spatial dimensions as the SRS images but with a dwell time of 8 µs pixel and averaged 6 times.

### Spontaneous Raman Microscopy

Spontaneous Raman spectra of target metabolites were acquired using a Horiba Xplora plus confocal Raman spectrometer equipped with a 532 nm YAG laser (12 mW) and a 100x, 0.9 N.A. objective (MPLAN N, Olympus) with a 500 µm pinhole and a 100 µm slit width. Spectra were collected with a 10 s integration time per frame and averaged over 10 frames using LabSpec6 software. Solvent background was subtracted from the metabolite spectra, followed by baseline correction and normalization.

## Acknowledgements

R.S.C. acknowledges financial support from the Biotechnology Leadership Pre-doctoral Training Program (BLP) Fellowship in the Donna and Benjamin M. Rosen Bioengineering Center at Caltech. A.C., J.A.A., and P.A.K. are grateful for financial support from a National Science Foundation Graduate Research Fellowship (DGE-1745301 and DGE-2139433, respectively). P.A.K. also acknowledges financial support from a Hertz Fellowship. L.W. is a Heritage Principal Investigator supported by the Heritage Medical Research Institute and acknowledges support from a CZI dynamic imaging grant. We also acknowledge helpful discussions with Noor Naji, Jacob-Parres Gold, Dr. Smruthi Karthikeyan, and Dr. Ryan Leighton.

## Author Contributions

R.S.C. and L.W. conceived the project. R.S.C., A.C., and L.W. designed the experiments. R.S.C. and A.C. conducted all experiments including SRS imaging. J.A.A. performed linear unmixing analysis. A.C. performed all quantitative analyses and generated the data visualizations. P.A.K. assisted in performing linear unmixing, data analysis, and data visualization. J.A.A and Z.Y. performed synthetic experiments. B.Y. assisted with data collection. The manuscript was written by R.S.C., A.C., and L.W. with input from all co-authors.

