## Supplementary Information for "MetaboRamics: Highly multiplexed metabolic imaging by stimulated Raman for spatial metabolomics in live cells"

### Supplementary Figures

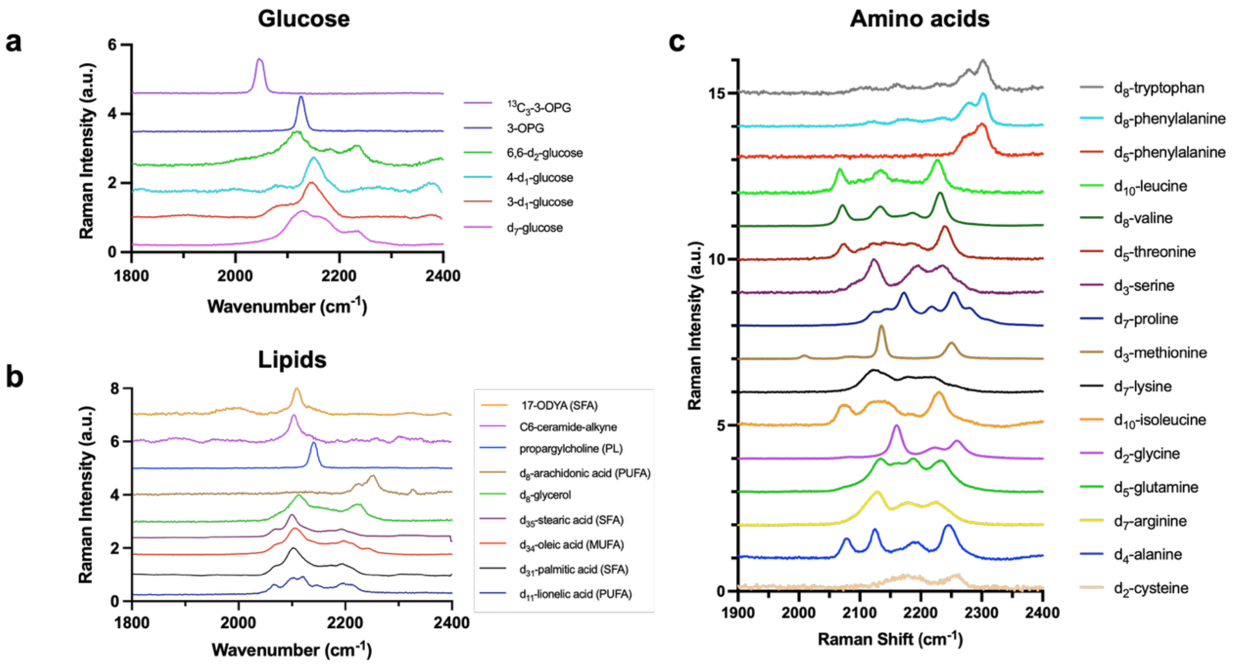

**Figure S1. Raman spectra of screened metabolites.** Spontaneous Raman spectra of different deuterium- or alkyne-based isotopologues or analogues of **a)** glucose; **b)** lipids and precursors; and **c)** amino acids; 3-OPG= 3-O-propargyl-d-glucose, 17-ODYA= 17-octadecynoic acid, SFA= saturated fatty acid, PL= phospholipids, MUFA= monounsaturated fatty acid, PUFA= polyunsaturated fatty acid.

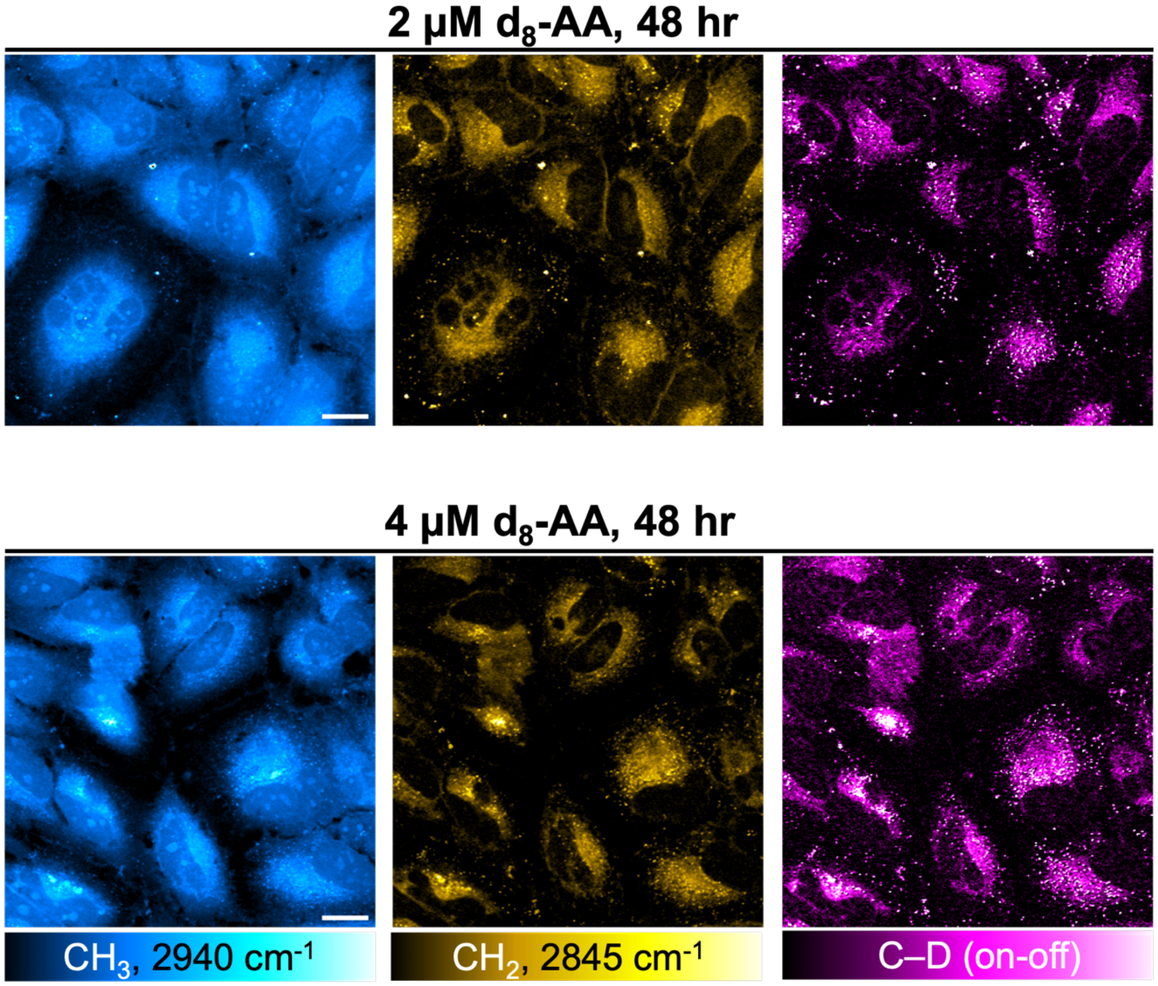

**Figure S2. Probe optimization for MetaboRamics.** Representative SRS images in live HeLa cells targeted at the protein ($CH_3$) and C–D (on-off) channel for cells incubated with only $d_8$-AA. We notice that an increase in incubation time (from 24 hr (condition used in **Fig. S5**) to 48 hr, top row) or concentration (from 2 μM to 4 μM, bottom row) increases the C–D signal intensity due to increased PUFA incorporation but also leads to formation of more LDs in cells.

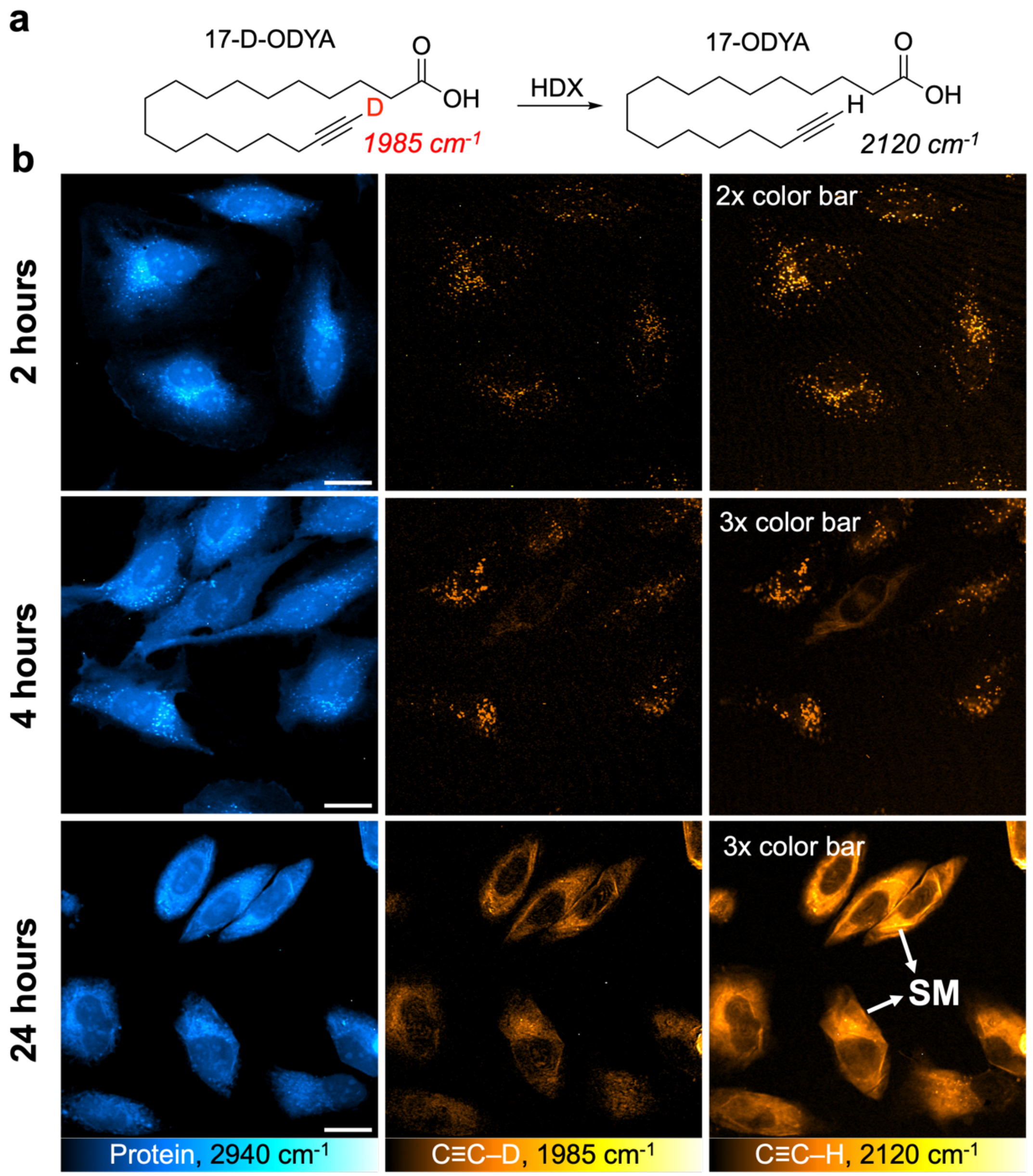

**Figure S3. HDX of 17-D-ODYA in live cells. a)** Chemical reaction indicating hydrogen-to-deuterium exchange (HDX) of 17-D-ODYA; **b)** Representative SRS images in live HeLa cells targeted at the protein ($CH_3$), C≡C–D (on-off) and C≡C–H (on-off) channels. Cells were incubated with 100 μM 17-D-ODYA for 2, 4, or 24 hr before SRS imaging. Patial HDX is evident at these incubation time points. Scale bar: 20 μm.

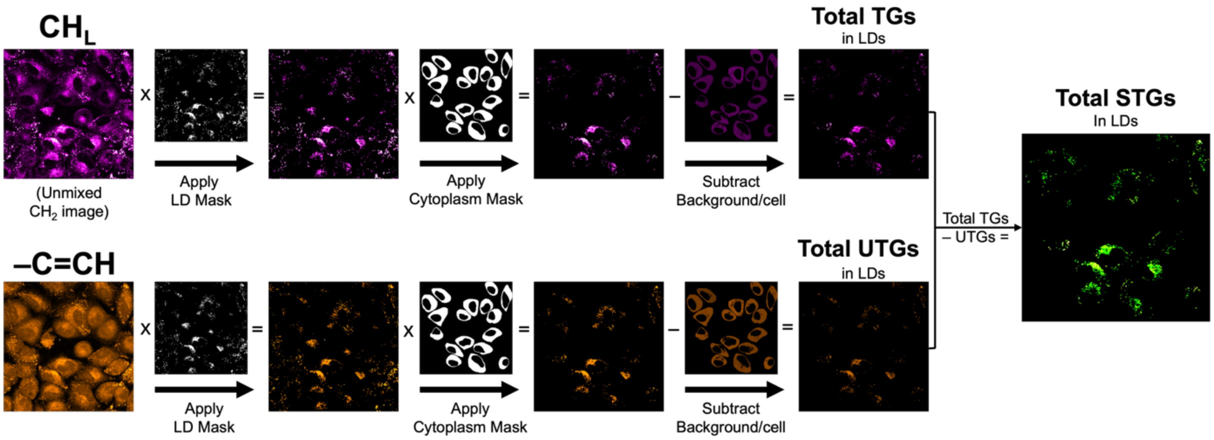

**Figure S4. Image processing workflow for calculation of saturated triglycerides (STGs) from SRS images.** STG images were prepared using a modified protocol adapted from a previous report.[1] Briefly, LDs were segmented to create binary LD masks from the CH$_L$ (unmixed $CH_2$ image, 2840 cm$^{-1}$) and UL (–C=C–H, 3022 cm$^{-1}$) images. These masks were then applied to their corresponding images. The resultant images for both CH$_L$ and U$_L$ were then multiplied with the cytoplasm mask to consider all LDs within the cell. Finally, the masked UL image was subtracted from the masked CH$_L$ image to yield the STG image (representing STGs obtained by subtracting unsaturated TGs from total TGs) in LDs.

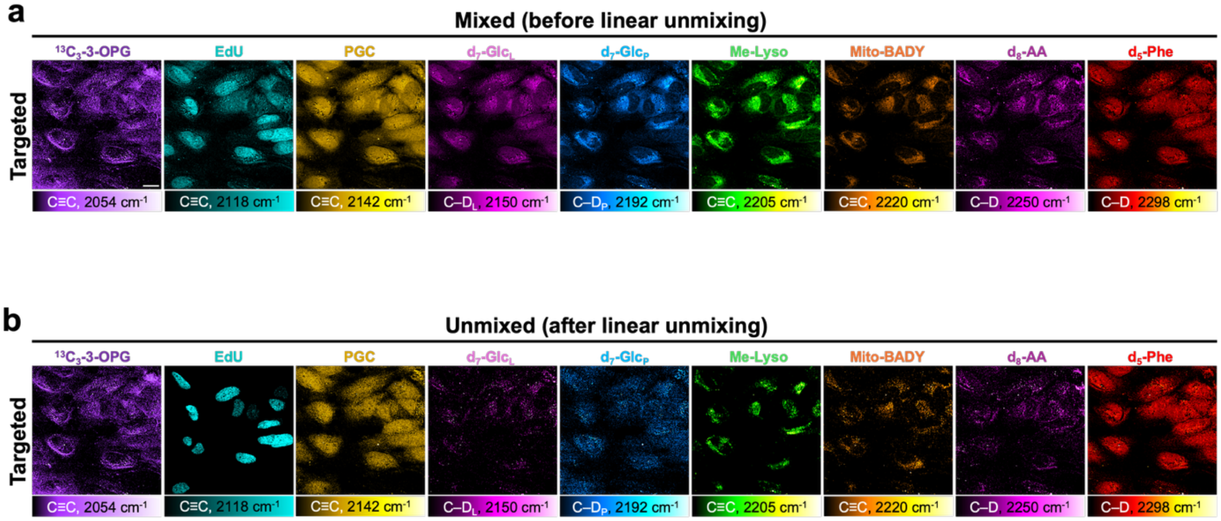

**Figure S5. Robust linear unmixing of targeted SRS channels.**
Comparison of **a)** mixed (before linear unmixing) and **b)** unmixed (after linear unmixing) SRS channels in live HeLa cells. Scale bar: 20 μm.

### Linear unmixing algorithm for 9 targeted SRS channels

Representative linear unmixing of the nine targeted SRS channels above with the exact recorded pump wavelength was performed using the algorithm provided below:

$$
\begin{bmatrix} I_{2053} \\ I_{2122} \\ I_{2142} \\ I_{2150} \\ I_{2192} \\ I_{2205} \\ I_{2221} \\ I_{2250} \\ I_{2298} \end{bmatrix}
=
\begin{bmatrix}
0.69 & 0.00 & 0.00 & 0.02 & 0.00 & 0.00 & 0.00 & 0.00 & 0.00 \\
0.02 & 0.90 & 0.05 & 0.63 & 0.58 & 0.00 & 0.00 & 0.00 & 0.00 \\
0.01 & 0.15 & 0.97 & 0.94 & 0.68 & 0.01 & 0.00 & 0.01 & 0.00 \\
0.01 & 0.04 & 0.58 & 1.00 & 0.48 & 0.01 & 0.00 & 0.01 & 0.00 \\
0.00 & 0.00 & 0.01 & 0.84 & 0.97 & 0.41 & 0.01 & 0.03 & 0.00 \\
0.00 & 0.00 & 0.01 & 0.69 & 0.88 & 1.00 & 0.07 & 0.06 & 0.00 \\
0.00 & 0.00 & 0.00 & 0.48 & 0.71 & 0.28 & 0.75 & 0.24 & 0.00 \\
0.00 & 0.00 & 0.00 & 0.26 & 0.33 & 0.01 & 0.08 & 0.77 & 0.03 \\
0.00 & 0.00 & 0.00 & 0.00 & 0.00 & 0.00 & 0.00 & 0.00 & 0.98
\end{bmatrix}
\times
\begin{bmatrix} \mathrm{C{\equiv}C}^{13}_{OPG} \\ \mathrm{C{\equiv}C}_{EdU} \\ \mathrm{C{\equiv}C}_{PGC} \\ \mathrm{CD}_{Lipid} \\ \mathrm{CD}_{Protein} \\ \mathrm{C{\equiv}C}_{Lyso} \\ \mathrm{C{\equiv}C}_{Mito} \\ \mathrm{CD}_{AA} \\ \mathrm{CD}_{Phe} \end{bmatrix}
$$

**inverse**

$$
\begin{bmatrix} \mathrm{C{\equiv}C}^{13}_{OPG} \\ \mathrm{C{\equiv}C}_{EdU} \\ \mathrm{C{\equiv}C}_{PGC} \\ \mathrm{CD}_{Lipid} \\ \mathrm{CD}_{Protein} \\ \mathrm{C{\equiv}C}_{Lyso} \\ \mathrm{C{\equiv}C}_{Mito} \\ \mathrm{CD}_{AA} \\ \mathrm{CD}_{Phe} \end{bmatrix}
=
\begin{bmatrix}
1.45 & 0.00 & 0.05 & -0.08 & 0.01 & 0.00 & 0.00 & 0.00 & 0.00 \\
-0.03 & 1.11 & 0.10 & -0.26 & -0.95 & 0.40 & -0.03 & 0.02 & 0.00 \\
0.00 & -0.22 & 1.55 & -0.85 & -0.83 & 0.34 & -0.02 & 0.00 & 0.00 \\
-0.02 & 0.15 & -1.59 & 2.65 & -0.43 & 0.17 & -0.01 & -0.01 & 0.00 \\
0.02 & -0.13 & 1.43 & -2.41 & 2.02 & -0.83 & 0.05 & -0.02 & 0.00 \\
0.00 & 0.01 & -0.16 & 0.26 & -1.37 & 1.60 & -0.13 & -0.03 & 0.00 \\
0.00 & 0.02 & -0.26 & 0.46 & -0.93 & -0.01 & 1.39 & -0.40 & 0.01 \\
0.00 & 0.00 & -0.05 & 0.08 & -0.61 & 0.28 & -0.16 & 1.35 & -0.04 \\
0.00 & 0.00 & 0.00 & 0.00 & 0.00 & 0.00 & 0.00 & 0.00 & 1.02
\end{bmatrix}
\times
\begin{bmatrix} I_{2053} \\ I_{2122} \\ I_{2142} \\ I_{2150} \\ I_{2192} \\ I_{2205} \\ I_{2221} \\ I_{2250} \\ I_{2298} \end{bmatrix}
$$

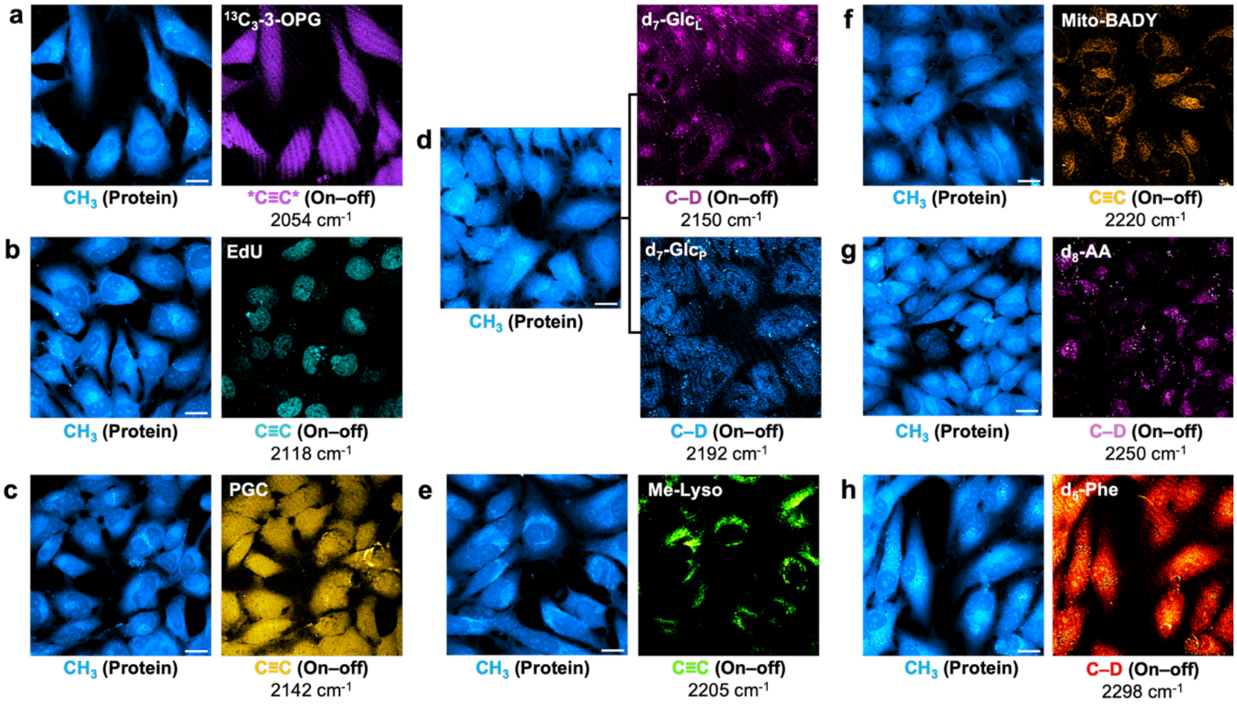

**Figure S6. Single-channel targeted SRS imaging. a-h)** Representative SRS images of optimized MetaboRamics metabolic probes (administered individually) alongside their corresponding protein ($CH_3$) images in live HeLa cells. As noted in the main text, $d_7$-Glc (in **d**) metabolizes into both lipids ($d_7$-$Glc_L$) and proteins ($d_7$-$Glc_P$), resulting in two spectrally distinct channels. Cells were treated using the same probe concentrations and incubation times as in the 16-plex MetaboRamics imaging experiments (see Materials & Methods). Scale bar: 20 μm.

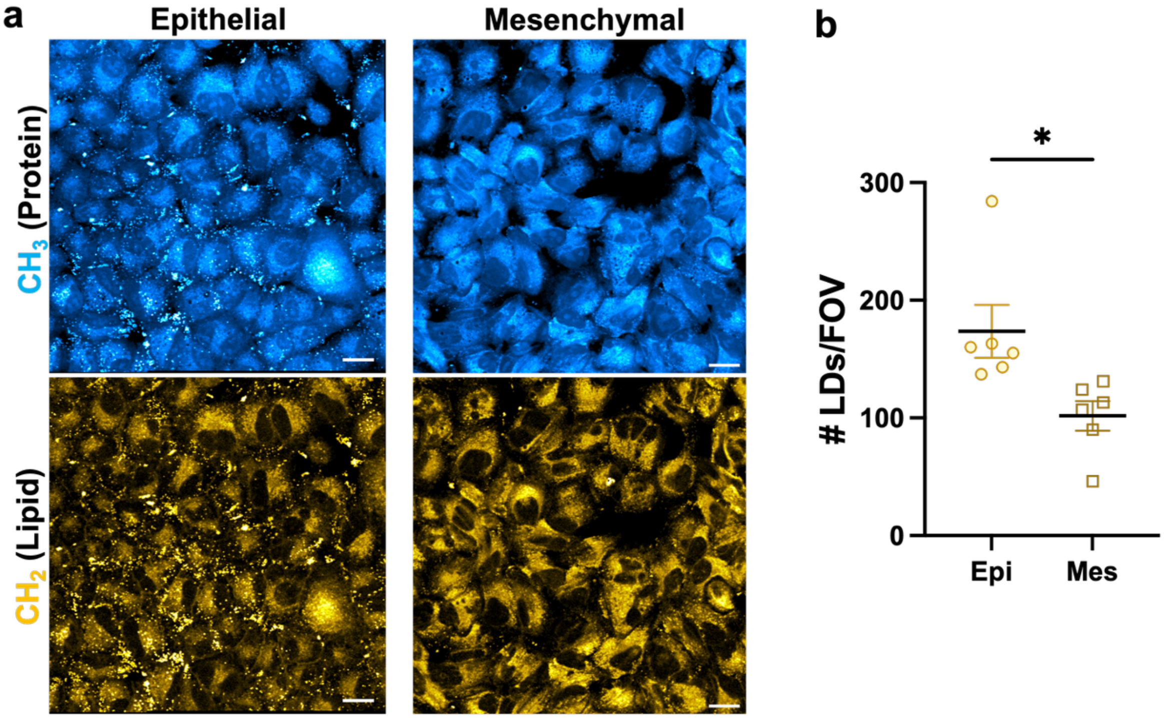

**Figure S7. Label-free assessment of lipid droplets (LDs) in EMT. (a)** Representative SRS images targeted at the protein ($CH_3$, top row) and lipid ($CH_2$, bottom row) channels for live A549 (epithelial and mesenchymal cells). Scale bar: 20 μm; **(b)** Quantification of number of LDs per FOV for epithelial and mesenchymal cells. Data represents mean (from different FOVs) ± SEM. Statistical significance was calculated using two-tailed Student's t-test, *$p<0.05$.

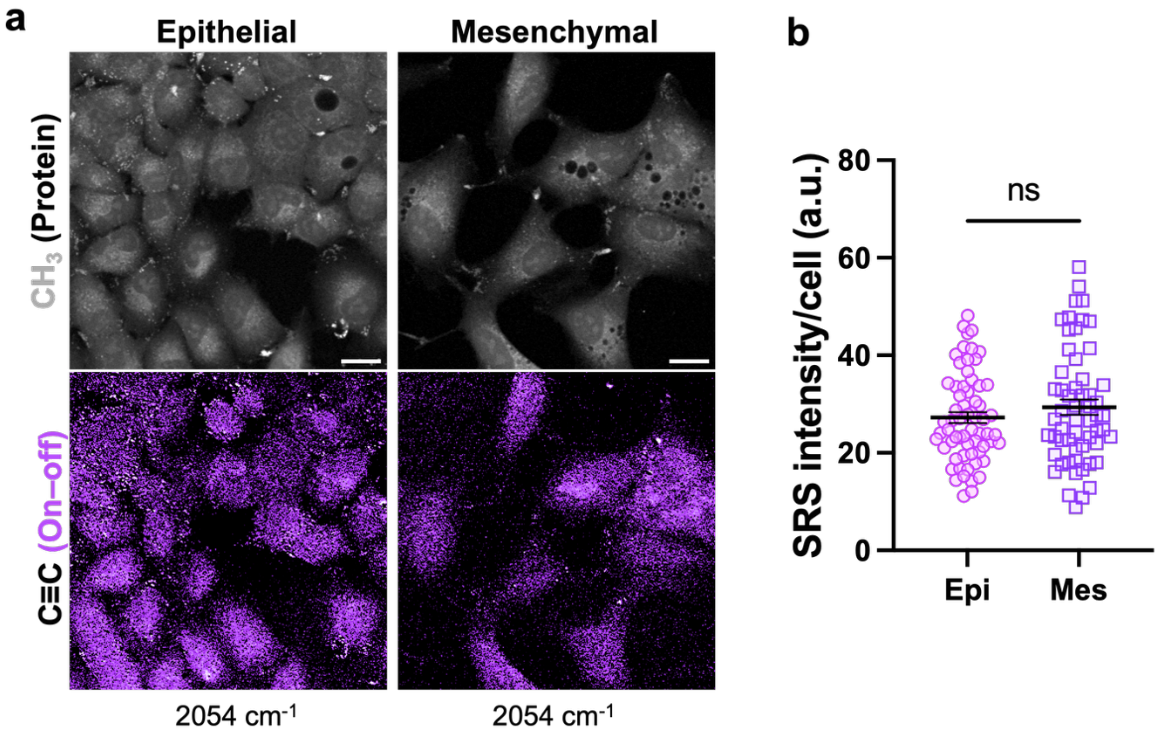

**Figure S8. Glucose uptake in EMT**. **(a)** Representative SRS images targeted at the protein ($CH_3$, top row) and alkyne (on–off) channels to monitor glucose uptake using 25 mM $^{13}C_3$-3-OPG in live epithelial and mesenchymal cells for 24 hours. Scale bar: 20 µm; **(b)** Quantification of SRS intensity per cell from alkyne images for epi and mes. Data represents mean ± SEM. Statistical significance was calculated using two-tailed Student's t-test, ns= no statistical difference.

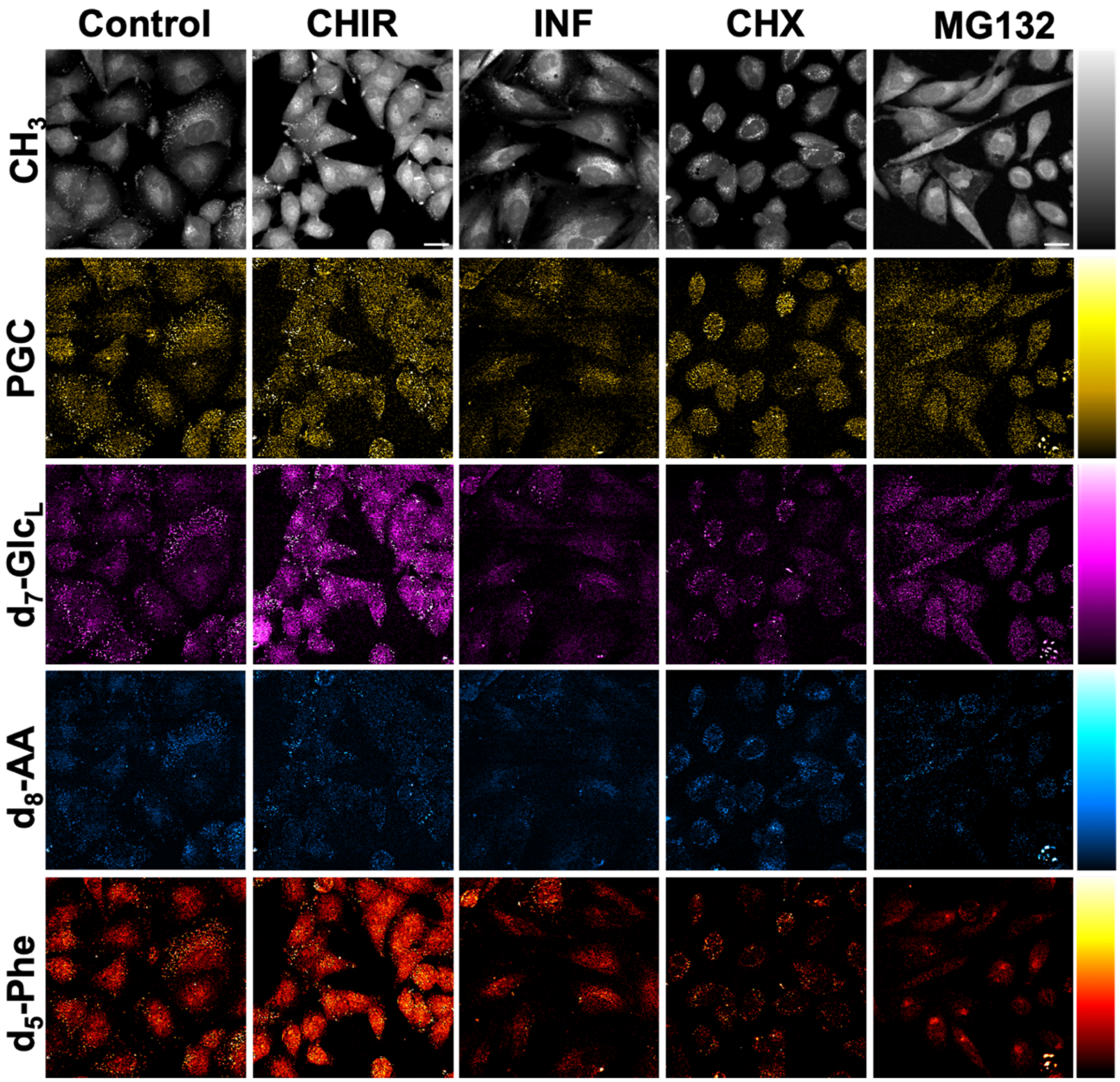

**Figure S9. SRS metabolic phenotyping of stress-induced cellular responses.** Select representative SRS images of live A549 cells under control conditions or treated with CHIR, INF, CHX, or MG132 with distinct metabolic signatures across stress states. Scale bar: 20 μm.

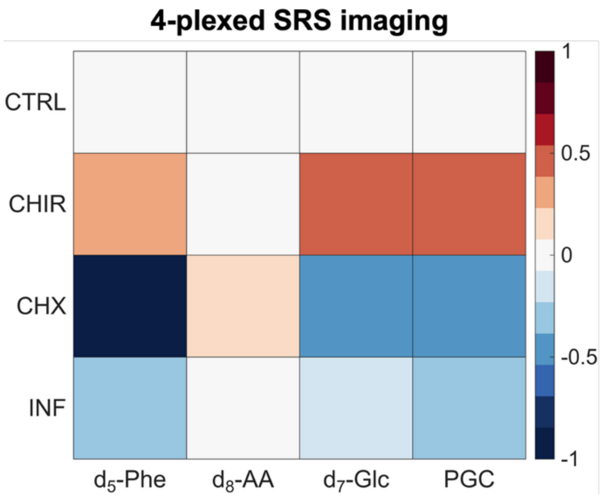

**Figure S10. Robustness of MetaboRamics.** Heatmap showing strong agreement between lower-order (4-plexed) and higher-order (16-plexed) imaging datasets, demonstrating the platform's scalability and reproducibility. Data from at least three independent experiments.

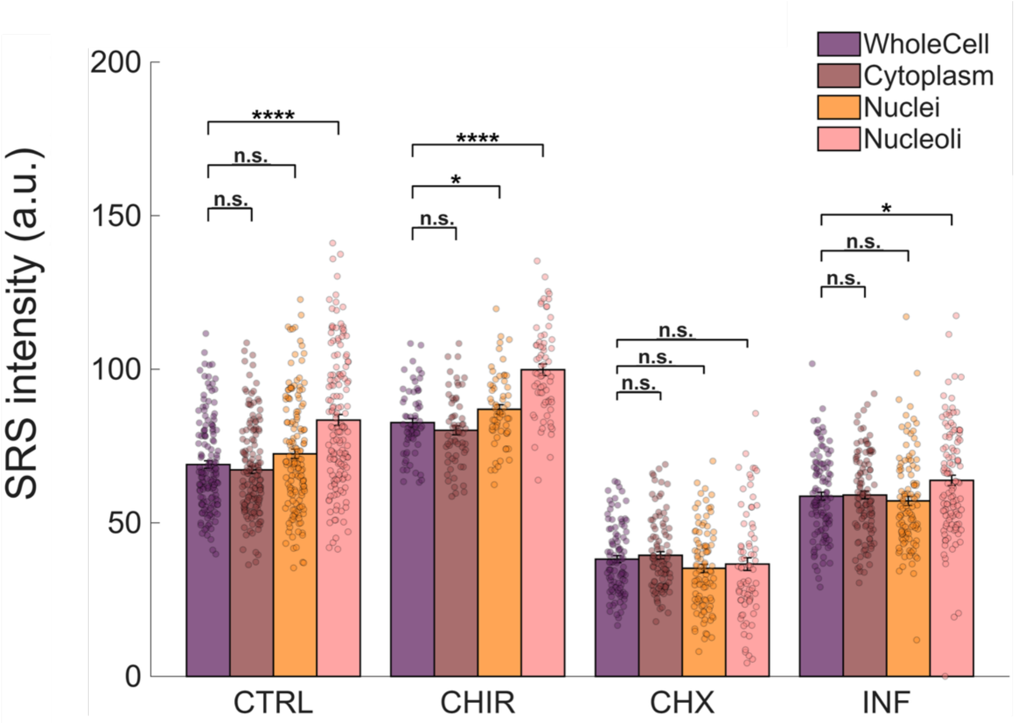

**Figure S11. Subcellular analysis of protein synthesis using MetaboRamics.** SRS intensities per cell for $d_5$-Phe at the subcellular level (whole-cell, cytoplasm, nuclei and nucleoli) for A549 control cells (no drug) or cells treated with INF or pharmacological perturbations: CHIR and CHX. Data represented as mean ± SEM from at least three independent experiments. Statistical analysis was conducted using Welch's two-paired t-test, *p<0.05, **p<0.01, ***p<0.001, ****p<0.0001 and n.s.= no statistical difference.

### Supplementary Materials and Methods

#### 1. Synthesis of MitoBADY (MB)

MitoBADY was synthesized following a protocol adapted from a previous report.[2] In a 100 mL flask, CuI (200 mg, 2 mmol) and TMEDA (600 μL, 4 mmol) were added to 10mL of acetone, and the mixture was bubbled with air for 10 minutes at room temperature. Next, 4-ethynylbenzyl alcohol (528 mg, 4 mmol, 1.0 eq), and phenylacetylene (1.32 mL, 12 mmol, 1.0 eq) were charged with 50 mL of degassed THF. The reaction was then stirred for 2 hours under air at room temperature. The mixture was concentrated in vacuo and subjected to flash column chromatography with a solvent gradient (0-1% MeOH in DCM), affording compound **1** in 35% yield.

In a 100 mL flame-dried flask, compound **1** is dissolved with 20 mL anhydrous DCM, and subsequently triethylamine (0.58 mL, 4.2 mmol, 1.2 eq) was added. The reaction mixture was then cooled with ice bath, and MsCl (0.17 mL, 2.09 mmol, 1.1 eq) was slowly injected into the reaction mixture, and the reaction was slowly raised to room temperature and allowed to stir at room temperature for 6 hours, and the reaction was monitored with TLC to observe almost full conversion after 6 hours. The reaction was quenched with 20 mL saturated sodium bicarbonate solution, and the reaction mixture was extracted with 50 mL DCM 3 times, and the combined organic phase was dried of $Na_2SO_4$, and the organic solvent was removed in vacuo, and the crude mixture was purified with flash column chromatography with DCM:MeOH=100:1 to afford the product compound with 42% yield.

In a 100 mL flame-dried flask, compound **2** (660 mg, 1.5 mmol, 1.0 eq) was added with $PPh_3$ (787 mg, 3.0 mmol, 2.0 eq) NaI (450 mg, 3.0 mmol, 2.0 eq) and dissolved with 25 mL anhydrous MeCN. The reaction was then stirred at 75°C for 6 hours, and the reaction was diluted with 100 mL DCM and filtered with celite, and the organic solvent was removed in vacuo, and the crude mixture was purified with flash column chromatography to afford the final product **3** with 77% yield.

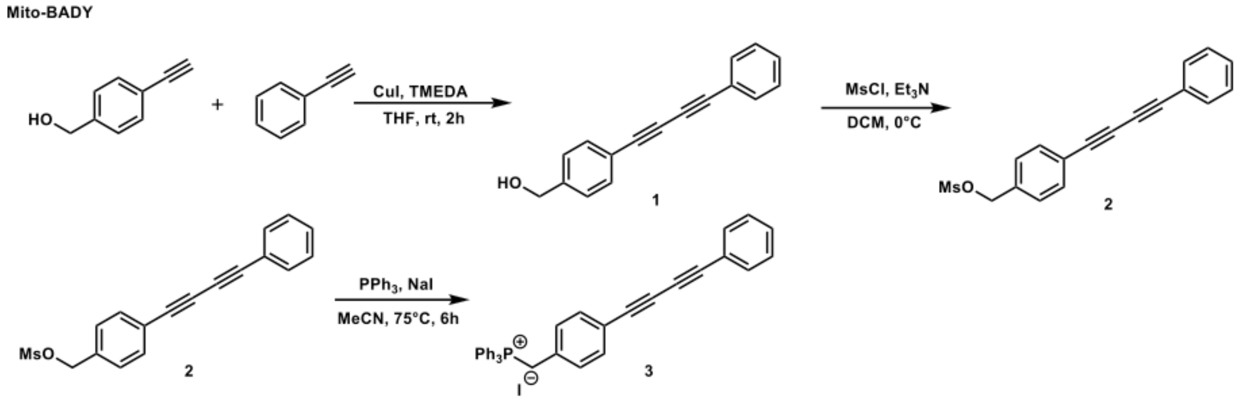

### 2. Synthesis of Me-Lyso (ML)

Me-Lyso was prepared using an improved adaptation of our previously reported protocol.[3] In a 250 mL flame-dried flask, 40 mL of anhydrous dichloromethane was charged with $AlCl_3$ (6.0 g, 44.8 mmol). The mixture was cooled to 0°C with an ice bath, followed by the dropwise addition of tetrachlorocyclopropene (2.0 g, 1.2 mL, 11.2 mmol). After 10 minutes of stirring at 0°C, mesitylene (1.344 g, 11.2 mmol) was added, and the reaction mixture was stirred for 90 minutes at 0°C. Next, 3,5-dimethylanisole (1.524 g, 11.2 mmol) was added, and the mixture was allowed to return to room temperature. After stirring for 1 hour, the mixture was lowered again to 0°C and quenched with $NH_4Cl$. The organic layer was extracted with dichloromethane, washed with brine, and dried over anhydrous $Na_2SO_4$. The crude mixture was purified with flash column chromatography with a solvent gradient (15-45% EA in hexanes), giving a mixture of Compounds **4** and **5** in 52% yield.

A 100 mL flame-dried flask was charged with the mixture of compounds **4** and **5** (1.74 g, 5.8 mmol) and 25 mL of anhydrous DCM. The mixture was lowered to 0°C, and 1M $BBr_3$ in DCM (15 mL, 15 mmol) was added slowly. The reaction mixture was raised to room temperature and allowed to run overnight. At 0°C, the reaction was slowly quenched with saturated $NH_4Cl$ solution. The organic phase was extracted with DCM, washed with brine, and dried over anhydrous $Na_2SO_4$. The crude mixture was purified with flash column chromatography with a solvent gradient (5-35% EA in hexanes), giving compounds **6** and **7** (4:1) in 22% yield.

In a 50 mL flame-dried flask was charged compound **6** (100 mg, 0.34 mmol, 1.0 eq) was added with (2-bromoethyl)dimethylamine hydrobromide(159 mg, 0.68 mmol, 2.0 eq) and $K_2CO_3$ (152 mg, 1.1 mmol, 3.5 eq). The mixture was dissolved with 10 mL dry DMF and the reaction was stirred at 70°C for 3 hours. The reaction was then quenched with 15 mL saturated $NH_4Cl$ aqueous solution and extracted with 30 mL ethyl acetate 3 times, the organic phase was dried over $Na_2SO_4$ and concentrated in vacuo. The crude mixture was then purified with flash column chromatography with Hex:EA=2:1 to afford the final product **8** as a white solid with 57% yield.

**Me-Lyso**

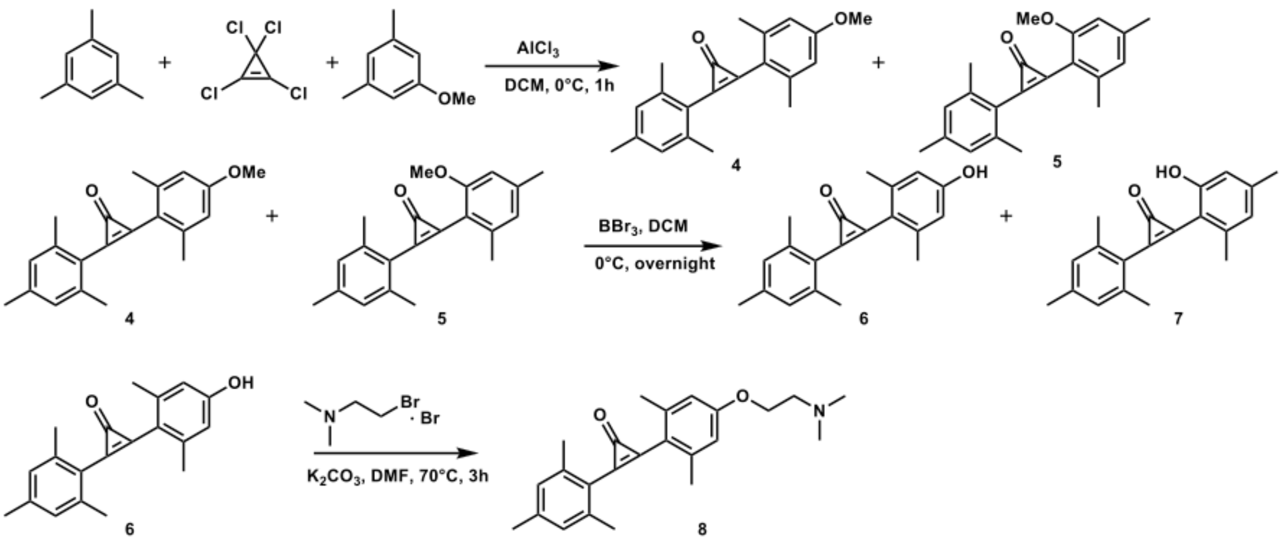
